# The lipid phosphatase-like protein PLPPR1 increases cell adhesion by modulating RhoA/Rac1 activity

**DOI:** 10.1101/470914

**Authors:** Chinyere Agbaegbu Iweka, Sharada Tilve, Caitlin Mencio, Yasuhiro Katagiri, Herbert M Geller

## Abstract

Phospholipid Phosphatase-Related Protein Type 1 (PLPPR1) is a member of a family of lipid phosphatase related proteins, integral membrane proteins characterized by six transmembrane domains. PLPPR1 is enriched in the brain and recent data indicate potential pleiotropic functions in several different contexts. An inherent ability of PLPPR1 is to induce membrane protrusions, and we have previously reported that members within this family may act in concert. However, the mechanism by which PLPPR1 produces these actions is not yet understood. Here, we report that exogenous expression of PLPPR1 reduces cell motility and increases cell adhesion to the ECM substrate by altering cytoskeletal dynamics and modulating RhoA and Rac1 activity through association with RhoGDI. This signaling also allows overexpression of PLPPR1 to overcome the inhibitory activity of CSPGs and LPA on neurites. Together, these results establish a novel signaling pathway for the PLPPR1 protein.

**SUMMARY:** PLPPR1 increases cell adhesion and decreases cell motility by modulating RhoA and Rac1 activation through its association with RhoGDI.

## INTRODUCTION

The Phospholipid Phosphatase Related (PLPPR) proteins (PLPPR1 – PLPPR5, previously referred to as Plasticity Related Genes, PRGs) are highly enriched in the brain and their expression is regulated, with the highest expression levels during development (Wang and Molnár, 2005). The first protein of this family to be discovered, PLPPR4, was found in sprouting axons following hippocampal deafferentation, implying a role in neuronal plasticity (Bräuer et al., 2003). Recent studies have shown that expression of PLPPR1 mRNA correlates with sprouting corticospinal axons after injury (Fink et al., 2017) and neuronal remodeling in the hippocampus after kainic acid treatment (Savaskan et al., 2004). Furthermore, decreased PLPPR1 mRNA has been associated with dysregulated neuronal migration (Khalaf-Nazzal et al., 2017; Pfurr et al., 2017), suggesting a role for these proteins in neuronal migration as well as neurogenesis and axon growth after injury.

We and others have previously shown that overexpression of PLPPR1 induces actin-rich membrane protrusions in many different cell types (Broggini et al., 2016; Sigal et al., 2007; Velmans et al., 2013; Yu et al., 2015). Overexpression of PLPPR5, the closest relative of PLPPR1 in this family, produces a similar phenotype (Broggini et al., 2010). The members of this family interact with each other and may function as a heteromeric complex to alter cytoskeletal dynamics (Yu et al., 2015). However, the mechanism by which PLPPR1 produces its effects is yet unknown.

Here we report that PLPPR1 expression leads to increased cell adhesion and changes in cytoskeletal dynamics that result in decreased cell migration. This enables cells overexpressing PLPPR1 to overcome the inhibitory activity of lysophosphatidic acid (LPA) and chondroitin sulfate proteoglycans (CSPGs). We show that these effects of PLPPR1 are due to alterations in the RhoA-ROCK signaling pathway mediated by PLPPR1 through association with RhoGDI. These results establish a novel signaling pathway for the PLPPR family of proteins.

## RESULTS

### PLPPR1 reduces cell migration and increases cell adhesion

Overexpression of PLPPR1 induces membrane protrusions in several different cell types (Broggini et al., 2016; Savaskan et al., 2004; Sigal et al., 2007; Velmans et al., 2013; Yu et al., 2015), implying a change in the cytoskeleton. Many cellular processes, including migration and adhesion, are governed by dynamic changes to the cytoskeleton. To assess the potential role of PLPPR1 on these processes, we expressed either EGFP or EGFP-PLPPR1 in Neuro2a cells, which do not express detectable levels of PLPPR1 mRNA (data from (Iida, 2018; Llorens et al., 2013) accessible at the NCBI GEO database), plated on a uniform fibronectin substrate. Live cell imaging was used to trace the migratory trajectories of individual cells (Fig. 1A). We found that the velocity (Fig. 1B) and total distance travelled (Fig. 1C) by migrating cells expressing PLPPR1 were significantly reduced compared to cells expressing EGFP.

**Figure 1.**
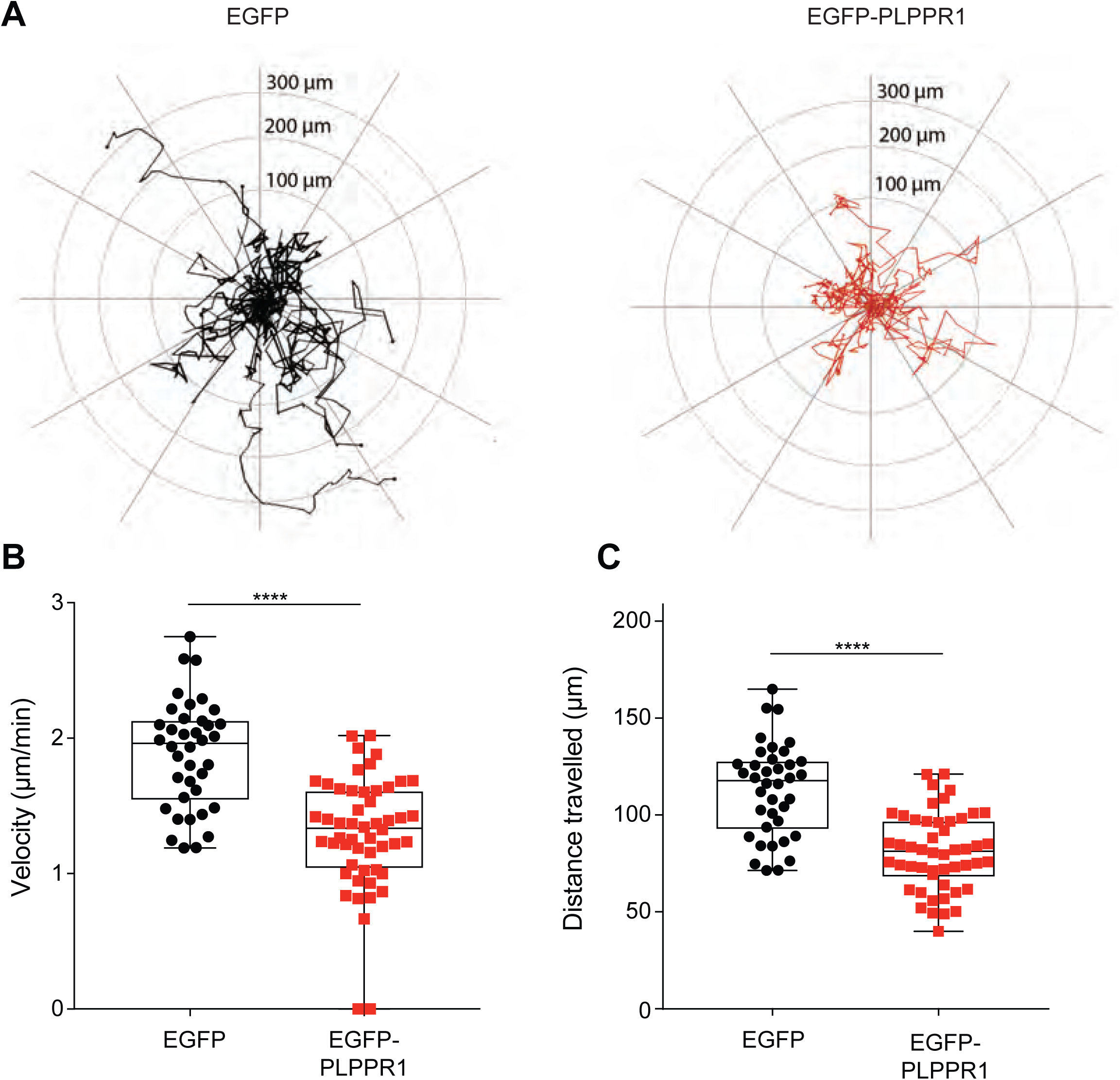
Cell migration is decreased in cells overexpressing PLPPR1. **(A)** Trajectories of Neuro2a cells transfected with pEGFP (left) or pEGFP-PLPPR1 (right), are plotted using the middle of each cell body as a point of reference. **(B)** Average velocity and **(C)** total distance travelled of migrating cells were determined using NIS Elements software. Data is represented as boxplot. EGFP (n = 38 cells; (B) Median 1.96, Range 1.19 to 2.75; (C) Median 117.7, Range 71.37 to 164.9); EGFP-PLPPR1 (n = 52 cells (B) Median 1.33 Range 0 to 2.02; (C) Median 81.32 Range 39.95 to 121.1). p-values were calculated using student’s *t*-test with Welch’s analysis, ****p<0.0001. Experiment was conducted in triplicate.

Cell adhesion is a major determinant for cell migration. We therefore performed both a cell adhesion and a cell detachment assay. Neuro2a cells expressing either EGFP or EGFP-PLPPR1 were plated on a fibronectin substrate. For the adhesion assay, cells were plated for one hour and subjected to media washes every 15 min, at which times the number of attached cells was measured. For the detachment assay, cells were allowed to attach overnight and their resistance to detachment by trypsin/EDTA was measured. Our results show that cells expressing PLPPR1 were both more adherent after media washing (Fig. 2A, B) and resistant to detachment by Trypsin/EDTA (Fig. S1).

**Figure 2.**
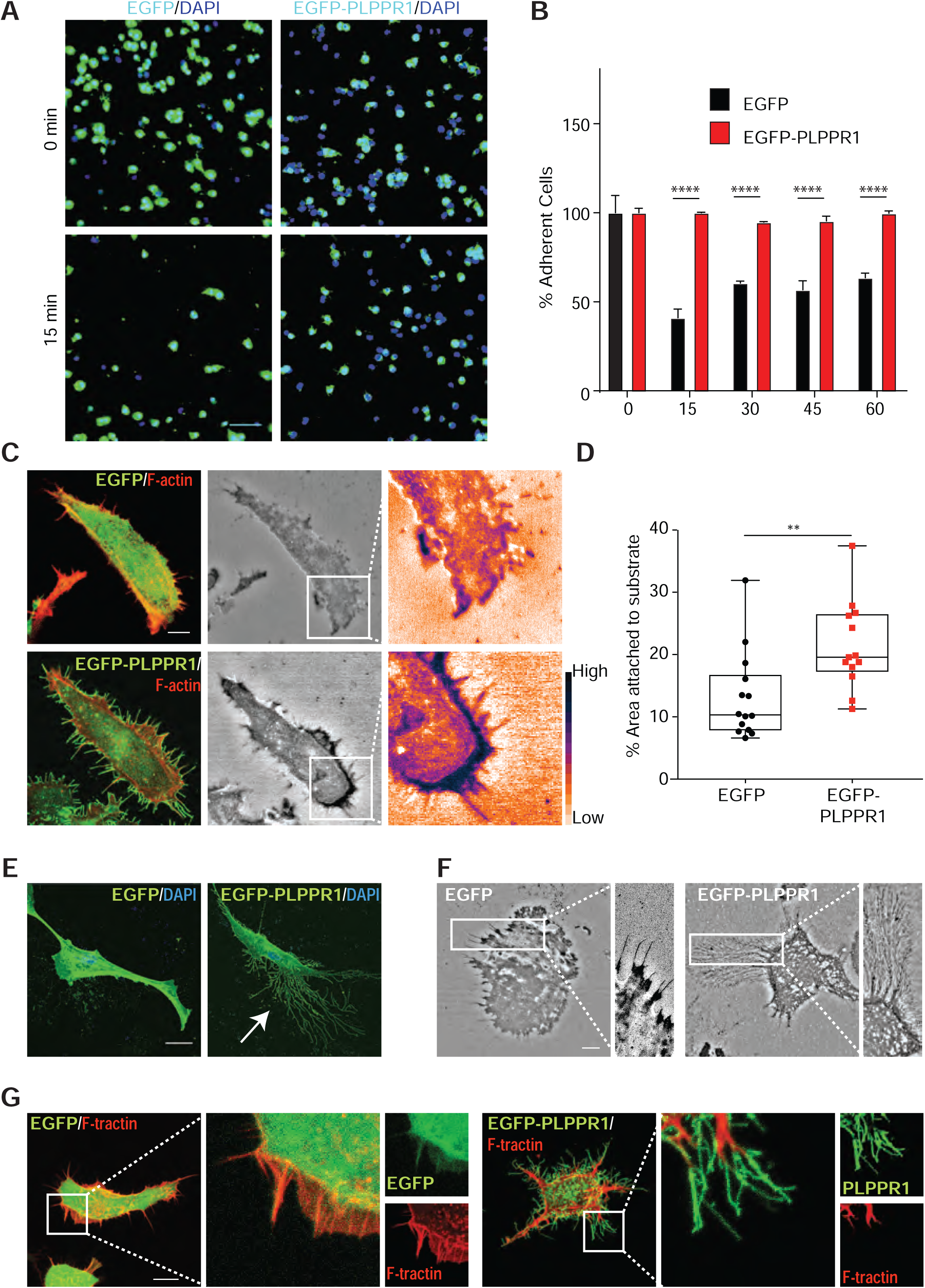
PLPPR1 increases cell adhesion. **(A, B)** Neuro2a cells expressing EGFP or EGFP-PLPPR1 were detached with 5 mM EDTA in PBS, re-plated on fibronectin coated 48-well plates and washed at 15, 30, 45 and 60 min. Scale bar, 100 µm. **(A)** Representative images at 15 min before (top) and after washing (bottom). **(B)** Quantification of the percentage of adherent cells after washing at different time points. Data represents mean ± SEM. Individual p-values were calculated using Two-way ANOVA with Tukey’s post-hoc analysis, ****p<0.0001. **(C)** Representative IRM images showing attachment of Neuro2a cells transfected with either pEGFP (top) or pEGFP-PLPPR1 (below) to the fibronectin substrate. Scale bar, 10 µm. Insets are higher magnification images of the boxed area. **(D)** Relative attachment of the cell to the fibronectin substrate was calculated by thresholding darker pixels divided by the traced cell area. An ImageJ color map ‘Gem’ is used to display higher intensity pixels as blue and lower intensity as orange which represent stronger and weaker adhesions respectively. Data is represented as boxplot (EGFP n = 14 cells; Median 19.57, Range 11.27 to 37.45) and (EGFP-PLPPR1, n = 13 cells; Median 10.31, Range 6.60 to 31.92) calculated using Student’s *t*-test with Welch’s analysis. **p<0.01. **(E)** Representative confocal images of cells overexpressing EGFP-PLPPR-1 cells (right) demonstrate trailing fibers (white arrow) which are absent in cells expressing EGFP (left). Nuclei are stained with DAPI (blue). Scale bar, 10 µm. **(F)** Representative IRM images of cells transfected with either pEGFP (left) or pEGFP-PLPPR1 (right). Inset is a higher magnification of boxed area showing trailing fibers in cells overexpressing PLPPR1. Scale bar, 10 µm. **(G)** Representative images of cells co-expressing F-tractin (mApple-FTR-940) with either EGFP (left) or EGFP-PLPPR1 (right) showing membrane protrusions are devoid of actin filaments. Scale bar, 10 µm. Inset is a higher magnification image of the boxed area. All experiments were conducted in triplicate.

To examine the intricacy of adhesion and mobility in cells expressing PLPPR1, we used Interference Reflection Microscopy (IRM) to visualize and quantify the interface between the ventral cell surface and the fibronectin substrate coated on glass. Cell membranes in close contact with the substrate produce more interference detected as darker pixels while membranes further from the substrate have less interference detected as lighter pixels (Fig. 2C). IRM showed that the average area of attachment to the fibronectin substrate was significantly greater in cells expressing PLPPR1, suggesting a stronger attachment compared to cells expressing EGFP (Fig. 2D). These observations confirm our results that show increased attachment in cells expressing PLPPR1.

Using live-cell IRM, we observed the formation of a trail of fibrous structures as PLPPR1-expressing cells migrate (Movie S1, Fig. 2E, F). These retraction fiber trails were formed in response to cell movement, as opposed to previously seen actin-rich protrusions, suggesting that these “trailing fibers” may be the result of the increased cell attachment. PLPPR1 overexpression is known to cause filopodia like actin-rich protrusions (Velmans et al., 2013; Yu et al., 2015). To determine the nature of these “trailing fibers”, we co-expressed mApple-FTR-940 (F-tractin, a cellular probe for filamentous actin) with either EGFP or EGFP-PLPPR1 in Neuro2a cells. Confocal images showed no expression of F-tractin in these “trailing fibers” (Fig. 2G). The presence of such “trailing fibers”, devoid of cytoskeletal components, have been attributed to active migration in cancer cells (DePasquale, 1998; Haemmerli and Strauli, 1981). This lack of actin implies they are not protrusions but may instead be remnants of cell membrane that, due to the PLPPR1-induced increase in cell adhesion, could not detach readily from the fibronectin substrate during migration.

### PLPPR1 increases nascent focal adhesion complexes

The increased adhesion observed in cells expressing PLPPR1 led us to take a closer look at focal adhesions (FA). Focal adhesions are mechanosensitive and require the recruitment of several proteins in order to change their composition and mature (Pasapera et al., 2010). This tension-mediated maturation involves the recruitment of paxillin to nascent focal adhesion complexes that are initially small in size, but then mature into focal adhesions as the size of the focal complexes increase and are stabilized (Pasapera et al., 2010; Zaidel-Bar et al., 2003). Therefore, cells expressing either EGFP or EGFP-PLPPR1 were plated on a fibronectin substrate, immunostained for paxillin and F-actin and imaging performed using TIRF (Fig. 3A). Measurement of the size of individual FA revealed more nascent paxillin in cells overexpressing PLPPR1 as compared to cells expressing EGFP, suggesting reduced FA turnover (Fig. 3B). Live cell imaging confirmed that focal adhesion complexes in cells co-expressing EGFP-PLPPR1 and mCherry-paxillin remained nascent compared to the mature FA observed in cells expressing EGFP (Movie S2).

**Figure 3.**
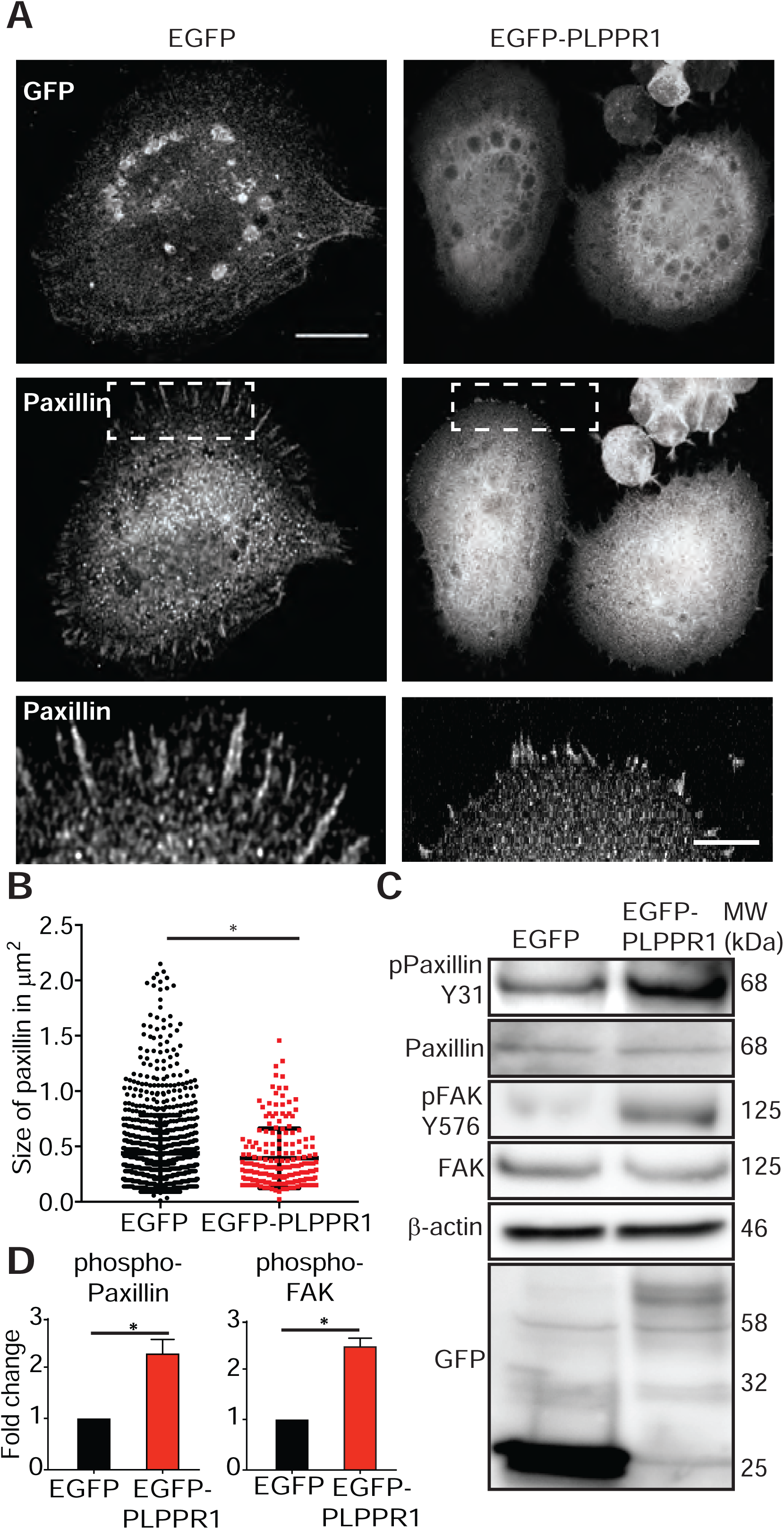
Focal adhesion contacts are increased in cells overexpressing PLPPR1. **(A)** Representative TIRF images of Neuro2a cells transfected with either pEGFP or pEGFP-PLPPR1, plated on fibronectin coated MatTek dishes and immunostained for paxillin. White dashed box area is presented at higher magnification of paxillin staining. Scale bar, 10µm and 1µm. **(B)** Mean size of paxillin were determined from FAs isolated with a segmentation algorithm in ImageJ. Data represents mean ± SEM. p-values were calculated using Student’s *t*-test with Welch’s analysis, *p<0.05, ****p<0.0001. **(C)** Immunoblot of transfected Neuro2a cells, probed for pY31, paxillin, pY576 and FAK. Membranes were stripped and reprobed for GFP and β-actin for loading control. **(D)** Densitometric analysis of phosphorylated protein versus total protein for paxillin and FAK was performed using Image Studio Lite. Data represents mean ± SEM. p-values were calculated using Student’s *t*-test with Welch’s analysis, *p<0.05. All experiments were performed in triplicate.

Phosphorylation of paxillin on Y31 is crucial for the regulation of paxillin turnover and FA complex maturation. Studies have demonstrated a reduced phospho-paxillin to total paxillin ratio in the transition from nascent to mature FA (Zaidel-Bar et al., 2007). Therefore, we assessed phosphorylation levels of paxillin to account for the nascent FAs observed in the PLPPR1-expressing cells. Immunoblotting of Neuro2A cell lysates revealed an increase in phosphorylation of paxillin in cells expressing PLPPR1 (Fig. 3C, D). Paxillin phosphorylation at Y31 is mediated by FAK (Bellis et al., 1995; Schaller and Parsons, 1995), therefore, we evaluated phosphorylation levels of FAK. Phosphorylation of FAK at Y576 in cells expressing PLPPR1 was increased compared to EGFP expressing cells (Fig. 3C, D). Altogether, these results suggest that the increased phosphorylation of paxillin and FAK in cells expressing PLPPR1 likely account for the increase in nascent FAs, resulting in increased cell adhesion (Beningo et al., 2001).

### PLPPR1 decreases the rate of actin polymerization

The dynamic organization of the actin cytoskeleton causes changes in cell shape and adhesion strength. This generates tension and creates the pushing and pulling forces that enable cell migration (Gardel et al., 2010; van Helvert et al., 2018). Given our results showing increased adhesion and decreased migration in cells expressing PLPPR1, we investigated the effect of PLPPR1 on the organization of the actin cytoskeleton. Confocal imaging demonstrated that Neuro2a cells expressing PLPPR1 lack the network of organized linear stress fibers observed in cells expressing EGFP (Fig. 4A). Because actin polymerization and depolymerization are required for stress fiber formation, we evaluated actin turnover by Fluorescence Recovery After Photobleaching (FRAP) measurements in cells co-expressing F-tractin with either EGFP or EGFP-PLPPR1 (Fig. 4B). The leading edge of the cell was photo-bleached and fluorescence recovery was measured. Results show a decrease in the rate of fluorescence recovery of actin in cells expressing PLPPR1 (Fig. 4C). The mobile fraction of monomeric actin was also decreased in PLPPR1-expressing cells (Fig. 4D).

**Figure 4.**
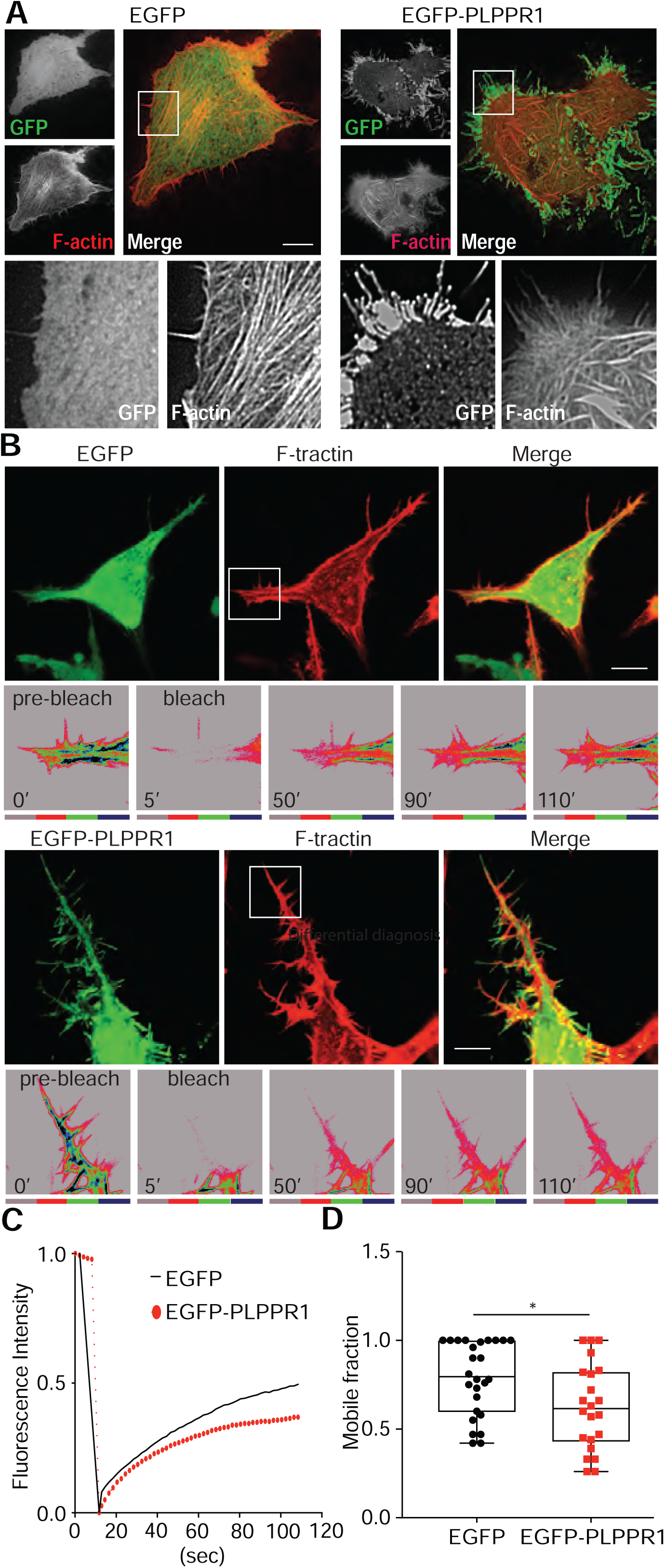
PLPPR1 expression induces changes in actin polymerization. **(A)** Actin stress fibers in Neuro2a cells expressing either EGFP (left) or EGFP-PLPPR1 (right) Scale bar, 10 µm. Inset is a higher magnification of boxed area. **(B)** FRAP analysis of actin polymerization in Neuro2a cells co-transfected with F-tractin (mApple-FTR-940) and either pEGFP (top) or pEGFP-PLPPR1 (bottom). Top images demonstrate cell morphology; the boxed areas were subjected to FRAP and imaged at the indicated times post bleaching. Scale bar, 10µm. **(C)** Graph shows fluorescence recovery over time in cells expressing EGFP (black solid line) and cells overexpressing PLPPR1 (red dotted line). **(D)** Mobile fraction was determined using Matlab. Data is represented as boxplot (EGFP Median 0.80, Range 0.42 to 1) and (PLPPR Median 0.62 Range 0.26 to 1). p-values were calculated using Student’s *t*-test with Welch’s analysis. *p<0.05.

### PLPPR1 overcomes CSPG and LPA neurite inhibition

Overexpression of PLPPR1 protein can overcome the inhibitory activity of LPA-induced axon collapse and neurite retraction (Broggini et al., 2016). Furthermore, PLPPR1 has been demonstrated to promote axon regeneration after spinal cord injury in mice (Fink et al., 2017). Chondroitin sulfate proteoglycans (CSPGs) are universally upregulated in the glial scar following injury and, similar to LPA, are potent negative regulators of axon growth (McKeon et al., 1991; Wang et al., 2008). Consequently, we asked if overexpression of PLPPR1 could overcome the inhibitory activity of CSPGs. Hippocampal neurons were transfected with EGFP or EGFP-PLPPR1 and seeded on glass coverslips coated with either poly-L-lysine (PLL) or PLL and CSPGs (Fig. 5A). Neurite outgrowth was analyzed by measuring total neurite length and the length of the longest neurite. Total neurite length of EGFP-transfected neurons was reduced by 37.5% in the presence of CSPGs. This reduction was not observed in PLPPR1-transfected neurons (Fig. 5B). Similarly, we observed a decrease in the length of the longest neurite of EGFP-transfected neurons, while this inhibition was not observed in neurons overexpressing PLPPR1 (Fig. 5C).

**Figure 5.**
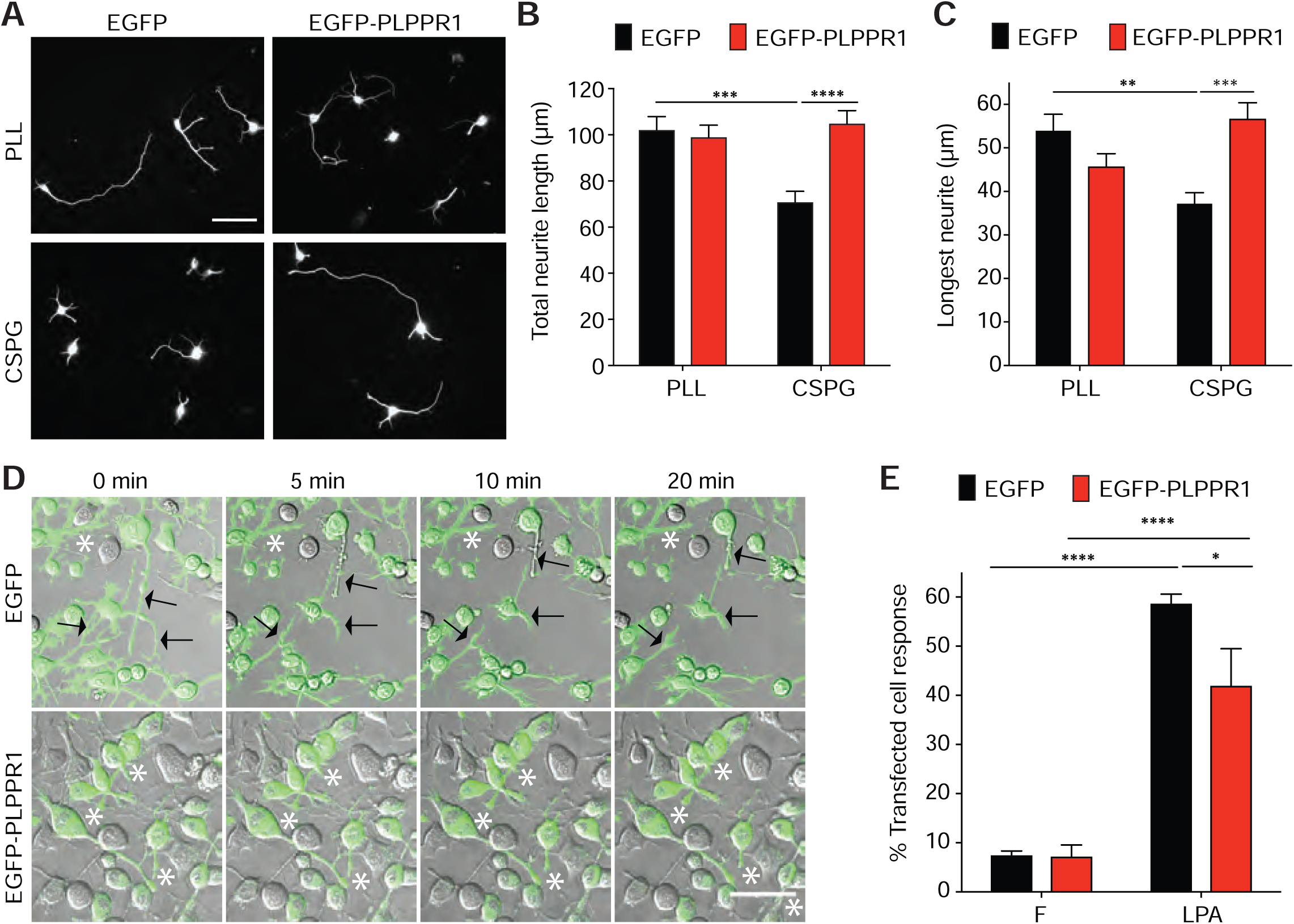
PLPPR1 attenuates LPA and CSPG mediated neurite inhibition and alters ROCK target phosphorylation. **(A)** Representative images showing pEGFP (left) or pEGFP-PLPPR1 (right) expressing hippocampal neurons cultured on either PLL (top) or PLL and CSPGs (bottom). Scale bar, 50 µm. **(B)** Analysis of total neurite length and **(C)** longest neurite were assessed at 2 DIV using NeuronJ. EGFP/PLL, n = 112; EGFP-PLPPR1/PLL, n = 115; EGFP/CSPGs, n = 113; EGFP-PLPPR1, n = 134 neurons. Data represent mean ± SEM. p-values were calculated using Two-way ANOVA with Tukey’s posthoc analysis, **p<0.01, ***p<0.001, ****p<0.0001. **(D)** Time lapse imaging of Neuro2a cells expressing either EGFP (top) or EGFP-PLPPR1 (bottom) and exposed to LPA. Black arrows indicate retracting neurites. White asterisks indicate nonretracting neurites. **(E)** Cell response was calculated as the percentage of transfected cells that retracted their neurites. Data represent mean ± SEM. p-values were calculated using Two-way ANOVA with Tukey’s posthoc analysis. *p<0.05, ****p<0.0001. Scale bar, 50µm. **(F)** Immunoblots of cells transfected with either pEGFP or pEGFP-PLPPR1 and treated with FAFBSA (F), serum (S), or LPA were probed for pMLC and MLC, pMYPT1 and MYPT1, pERM and ERM and **(G)** GFP. Densitometry analysis of phosphorylated MLC **(H)**, MYPT1 **(I)**, and ERM **(J)** versus total protein was performed using Image Studio Lite. Data represents mean ± SEM. p-values were calculated using Two-way ANOVA with Tukey’s posthoc analysis, *p<0.05, **p<0.01, ***p<0.001, ****p<0.0001 respectively. All experiments were conducted in triplicate.

LPA induces process retraction and cell rounding in several different cell types (Jalink et al., 1993; Kranenburg et al., 1999; Tigyi et al., 1996). To confirm this in Neuro2a cells, serum starved cells were exposed to either fatty acid free bovine serum albumin (FAFBSA) or LPA. Live cell imaging revealed significant neurite retraction and cell rounding in cells exposed to LPA compared to FAFBSA (Movie S3). Next, the effects of PLPPR1 expression on neurite retraction in Neuro2A cells induced by LPA was examined. After transfection with pEGFP or pEGFP-PLPPR1, cells were exposed to either FAFBSA or LPA. PLPPR1 expression reduced the response to LPA: 59% of cells expressing EGFP underwent cell rounding and neurite retraction in response to LPA compared to 42% of cells expressing PLPPR1 (Fig. 5D, E; Movie S4). Together, these results show that PLPPR1 expression impedes the inhibitory activity of CSPGs on neurite growth and attenuates LPA-induced neurite retraction and cell rounding.

### PLPPR1 modulates RhoA activation

The RhoA signaling pathway is common to many molecules inhibitory to axon growth, including CSPGs and LPA (Alabed et al., 2006; Jain et al., 2004; Kranenburg et al., 1999; Monnier et al., 2003). Additionally, RhoA activation by LPA is reduced in cells overexpressing PLPPR5, the closest relative to PLPPR1 (Broggini et al., 2010). Therefore, we asked if PLPPR1 could similarly affect RhoA activation by LPA. Serum starved Neuro2a cells expressing EGFP or EGFP-PLPPR1 were treated with either FAFBSA, serum or LPA. GTP-bound RhoA was immunoprecipitated using rhotekin-RBD beads and then subjected to immunoblot analysis. While cells expressing EGFP had increased RhoA activation in response to either serum or LPA, RhoA activation remained unchanged after serum or LPA treatment in cells expressing PLPPR1 (Fig. 6A, B).

**Figure 6.**
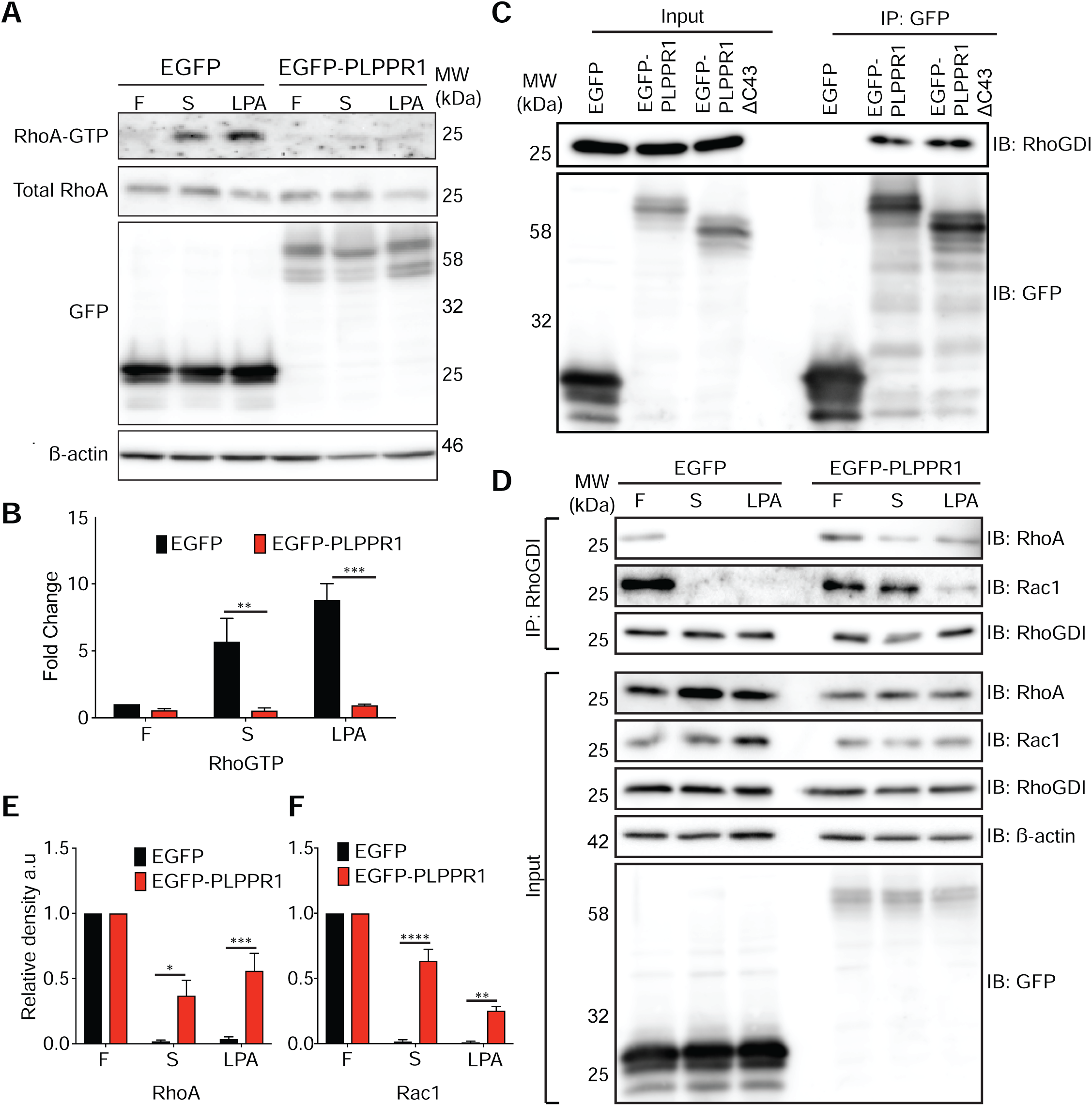
PLPPR1 reduces RhoA activation and maintains RhoA and Rac1 interaction with RhoGDI after LPA treatment. **(A)** RhoA-GTP was pulled down using rhotekin-RBD beads from cell lysates prepared from transfected Neuro2a cells treated with either FAFBSA (F), 10% serum (S) or 16 µM LPA for 2 min. Immunoblots were probed for RhoA. β-actin was used as loading control for total RhoA. Membranes were stripped and reprobed for GFP. **(B)** Densitometric analysis was performed using Image Studio Lite. Data represent mean ± SEM. p-values were calculated using Two-way ANOVA with Tukey’s posthoc analysis, **p<0.01, ***p<0.001. All experiments were performed in triplicate. **(C)** Neuro2a cells expressing either EGFP, EGFP-PLPPR1 or EGFP-PLPPR1ΔC43 were subjected to immunoprecipitation with GFP-MA^TM^ beads and immunoblotted with anti-RhoGDI antibody. Input was probed for GFP. **(D)** Serum starved transfected Neuro2a cells were treated with either FAFBSA (F), serum (S) or LPA and subjected to immunoprecipitation with RhoGDI antibody. Immunoblots of input and immunoprecipitates were probed for either RhoGDI, RhoA or Rac1. Inputs were stripped and reprobed for GFP and β-actin antibody for loading control. Densitometric analysis of RhoA **(E)** and Rac1 **(F)** was performed using Image Studio Lite. Data represents mean ± SEM. p-values were calculated using Two-way ANOVA with Tukey’s posthoc analysis, *p<0.05, **p<0.01, ***p<0.001, ****p<0.0001 respectively. All experiments were performed in triplicate.

Rho-associated kinase (ROCK) is a major RhoA effector and is implicated in the regulation of actomyosin dynamics and inhibition of neurite outgrowth (Amano et al., 1997; Lingor et al., 2007; Monnier et al., 2003). ROCK mediates its effect by phosphorylating several substrates including cytoskeletal proteins (Kaneko-Kawano et al., 2012; Matsui et al., 1998; Riddick et al., 2008; Sutherland et al., 2016; Yoneda et al., 2005). We investigated the effect of PLPPR1 on the RhoA-ROCK pathway by assessing the phosphorylation levels of the ROCK substrates MLC, MYPT1, and ERM in response to LPA treatment. Cell lysates were collected from serum starved Neuro2a cells expressing EGFP or EGFP-PLPPR1 that were treated with either FAFBSA, serum or LPA and immunoblotted for levels of pMLC, pMYPT1, and pERM. Densitometry analysis demonstrated an increase in phosphorylation of S19 MLC, T853 MYPT1 and T567, T564, T558 ERM proteins in EGFP-expressing cells in response to increasing concentrations of LPA, with no change in the phosphorylated levels in cells expressing PLPPR1 (Fig S2A). GFP immunoblots showed that expression levels for both EGFP and EGFP-PLPPR1 do not change between treatment conditions (Fig. S2B).

LPA is also known to induce changes in phosphorylation of Akt, GSK3ß and Tau through activation of the PI3K-PKB/Akt and PLCγ-PKC pathways respectively (Baudhuin et al., 2002; Fang et al., 2002; Sayas et al., 2006). Therefore, we determined if the effects of PLPPR1 expression extend to these pathways. Immunoblot analysis showed no significant change in phosphorylation levels of GSK3ß (Y216 and S9), Tau (S396) and Akt (S473) in cells expressing PLPPR1 compared to cells expressing EGFP after treatment with LPA (Fig. S3A).

To determine if PLPPR1 had any effect on the phosphorylation levels of ROCK target proteins without LPA stimulation, we performed an immunoblot analysis of MLC, ERM and MYPT1, including Akt, GSK3β or Tau, using cell lysates from Neuro2a cells expressing EGFP or EGFP-PLPPR1. Results show no significant changes in phosphorylation levels of these target proteins in cells expressing PLPPR1 compared to cells expressing EGFP (Fig. S3B). Altogether, these results demonstrate that PLPPR1 specifically attenuates activation of the RhoA-ROCK signaling pathway in response to stimulation.

### PLPPR1 modulates RhoA and Rac1 activation via RhoGDI interaction

Previously, we identified RhoGDI as a possible binding partner for PLPPR1 (Yu et al., 2013). By direct interaction with the Rho GTPases, RhoGDI regulates their translocation from the cytosol to the plasma membrane and impedes the nucleotide exchange of GDP for GTP (Boulter et al., 2010; Dovas and Couchman, 2005; Sabbatini and Williams, 2013). To confirm the interaction between RhoGDI and PLPPR1, we immunoprecipitated EGFP using GFP magnetic/agarose beads from cells expressing either EGFP or EGFP-PLPPR1 and immunoblotted for RhoGDI. The PLPPR family share high sequence homology except for the C-terminus, which is unique to each family member. Therefore, we also expressed EGFP-PLPPR1ΔC43 to determine if this putative interaction is dependent on the C-terminal domain. Our results show that both EGFP-PLPPR1 and EGFP-PLPPR1ΔC43 co-immunoprecipitated with RhoGDI, confirming that PLPPR1 associates with RhoGDI and this association is independent of the C-terminus (Fig. 6C).

Inactive RhoA (GDP bound form) is tightly sequestered by RhoGDI in the cytosol (Garcia-Mata et al., 2011; Michaelson et al., 2001). LPA stimulation results in the displacement of RhoA from RhoGDI, which allows RhoA to disengage from GDP, bind GTP and become active (Kranenburg et al., 1997). To determine if expression of PLPPR1 would affect the RhoGDI – RhoA interaction after LPA treatment, Neuro2a cells transfected with EGFP or EGFP-PLPPR1 were treated with either FAFBSA, serum or LPA. RhoGDI was immunoprecipitated and immunoblot analysis of RhoA was performed. As expected, after treatment with either serum or LPA, the interaction between RhoA and RhoGDI was disrupted in cells expressing EGFP. However, the affinity of RhoGDI for RhoA was maintained in cells expressing PLPPR1 even after serum or LPA treatment (Fig. 6D, E). Immunoblot analysis of GFP expression showed that protein expression levels did not change between treatment conditions (Fig. 6D).

Like RhoA, Rac1 activation is regulated by RhoGDI (Boulter et al., 2010; DerMardirossian et al., 2004). We assessed the effects of PLPPR1 expression on the affinity of RhoGDI for Rac1 after LPA treatment. Neuro2a cells expressing EGFP or EGFP-PLPPR1 were exposed to either FAFBSA, serum or LPA. RhoGDI was immunoprecipitated and immunoblotting was performed to detect Rac1 levels. Our results show that, like RhoA, the interaction between RhoGDI and Rac1 was disrupted by LPA treatment in cells expressing EGFP, however, this interaction was not disrupted in cells expressing PLPPR1 (Fig. 6D, F). Together, these results demonstrate that PLPPR1 modulates LPA induced activation of RhoA and Rac1, likely via its association with RhoGDI.

### Constitutively active (CA) Rac1 overexpression rescues PLPPR1 induced morphological change

Rac1 activation is essential to lamellipodial formation (Ehrlich et al., 2002; Kozma et al., 1997; Somanath and Byzova, 2009). Given the possibility that PLPPR1 may regulate Rac1 activity, we examined the morphology of cells expressing dominant negative (DN) Rac1 T17N, constitutively active (CA) Rac1 G12V and cells co-expressing PLPPR1 with CA Rac1. We observed that expression of DN Rac1 alone led to the formation of “trailing fibers” similar to those observed in cells expressing PLPPR1 (Fig. 7A). Interestingly, cells co-expressing CA Rac1 with PLPPR1 had fewer trailing fibers (Fig. 7A). We quantified the percentage of cells that displayed trailing fibers and saw a significant reduction in this phenotype when co-expressing CA Rac1 with PLPPR1 as compared to PLPPR1 alone (Fig 7B).

**Figure 7.**
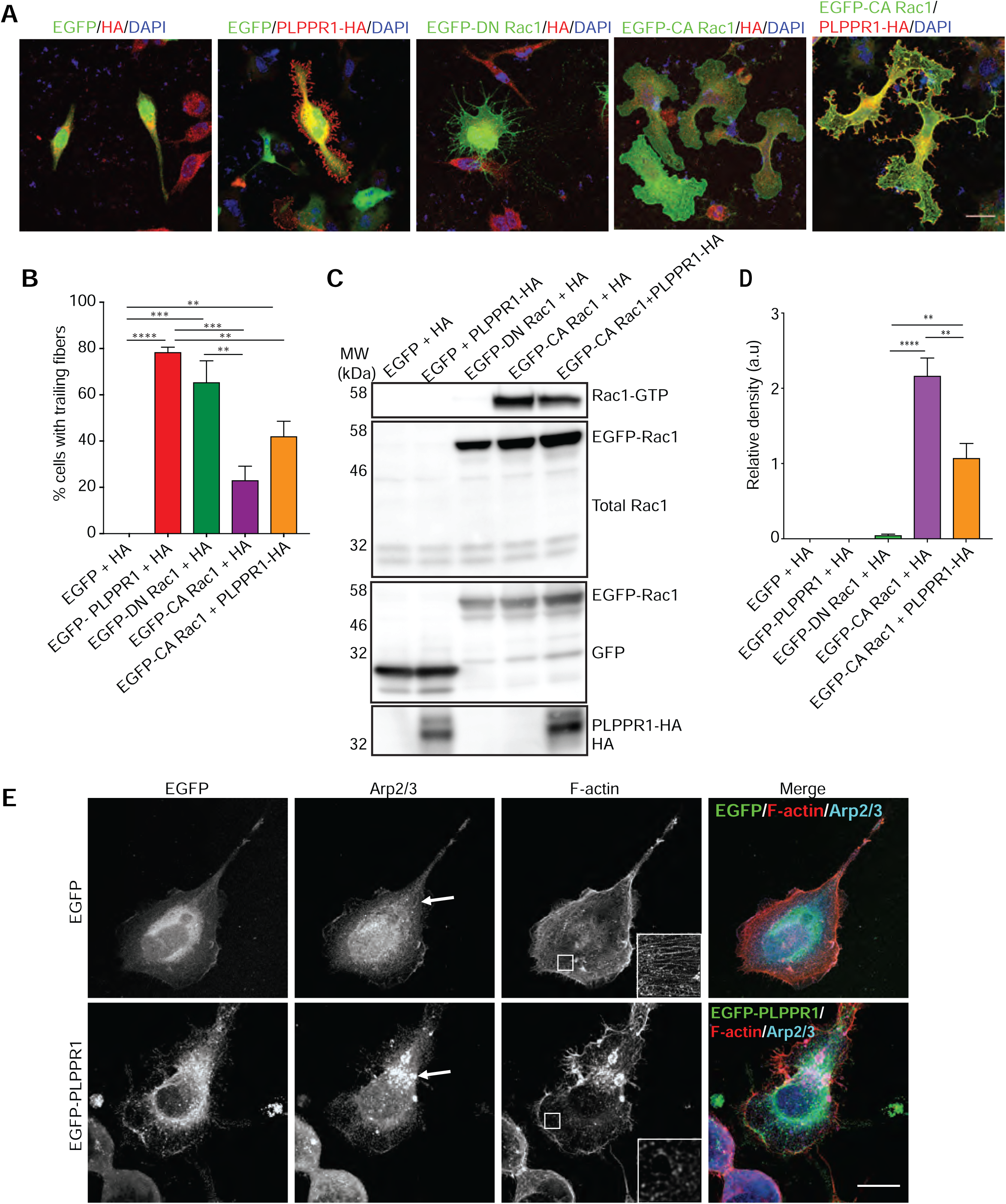
Co-expression of constitutively active Rac1 with PLPPR1 rescues morphological change. **(A)** Representative confocal images of Neuro2a cells expressing EGFP+HA, EGFP+PLPPR1-HA, EGFP-DN Rac1+HA, EGFP-CA Rac1+HA and EGFP-CA Rac1+PLPPR1-HA. Cells expressing EGFP-CA Rac1+PLPPR1-HA show a rescue of the morphological change induced by expression of PLPPR1. Scale bar, 10 µm. **(B)** Percentage of cells expressing the PLPPR1 trailing fibers phenotype of Neuro2a cells expressing EGFP+HA, EGFP-PLPPR1+HA, EGFP-DN Rac1+HA, EGFP-CA Rac1+HA and EGFP-CA Rac1+PLPPR1-HA**. (C)** Rac1-GTP was pulled down using PAK-PBD beads from cell lysates prepared from Neuro2a cells expressing EGFP+HA, EGFP+PLPPR1-HA, EGFP-DN Rac1+HA, EGFP-CA Rac1+HA and EGFP-CA Rac1+PLPPR1-HA. Membranes were immunoblotted with Rac1 antibody. Membranes were stripped and reprobed with anti-GFP antibody. **(D)** Densitometric analysis of Rac GTP versus total Rac1 was performed using Image Studio Lite. Data represents mean ± SEM. p-values were calculated using One-way ANOVA with Tukey’s posthoc analysis, **p<0.01, ****p<0.0001. **(E)** Cells transfected with either pEGFP (top) or pEGFP-PLPPR1 (bottom) were immunostained for Arp2/3, and phalloidin to visualize F-actin. White arrows indicate localization of Arp2/3. Inset is a higher magnification of boxed area. Scale bar, 10 µm. All experiments were conducted in triplicate.

To further explore the effect of PLPPR1 on Rac1 activation, DN Rac1 and CA Rac1 were expressed in Neuro2a cells and active Rac1 bound to GTP was pulled down using PAK-PBD beads. Basal levels of Rac1-GTP were near zero in Neuro2a cells expressing EGFP and DN Rac1, as well as in cells expressing PLPPR1 (Fig. 7C, D). High levels of active Rac1 were detected in cells expressing CA Rac1, as expected. Interestingly, co-expression of CA Rac1 with PLPPR1 significantly reduced active Rac1 (Fig. 7C, D).

Arp2/3 is an actin nucleating complex that acts downstream of Rac1 and is essential to actin assembly and lamellipodia formation (Lai et al., 2008; Machesky and Hall, 1997; Machesky and Insall, 1998; Wu et al., 2012). We examined Arp2/3 in cells expressing PLPPR1 using STED microscopy. Results show that Arp2/3 was localized to the actin-rich lamellipodia and exhibited a diffuse cytoplasmic distribution in cells transfected with pEGFP. Cells expressing PLPPR1 displayed a compact aggregation of Arp2/3 (Fig. 7E), suggesting a disruption in Arp2/3 localization. Together, these results suggest a modulatory effect of PLPPR1 on Rac1 activity, possibly through increased binding and extraction of active Rac1 by RhoGDI.

## DISCUSSION

The PLPPRs family has five members, of which PLPPR1 and PLPPR5 are most closely related. Overexpression of either PLPPR1 or PLPPR5 in cultured cells leads to a morphological change, including a significant increase in cellular protrusions in many cell types (Broggini et al., 2016; Velmans et al., 2013; Yu et al., 2015). Recent studies using genetic screening approaches have identified changes in PLPPR1 mRNA in models that show alterations of neurogenesis (Pfurr et al., 2017), neuronal migration (Khalaf-Nazzal et al., 2017), regulation of axonal growth (Fink et al., 2017), as well as in disease models such as schizophrenia (Windrem et al., 2017). However, the mechanism of action by which PLPPR1 induces morphological changes in cells and alters neural development has been a subject of speculation. We now show that PLPPR1 expression, through modulation of GTPase signaling, alters cell adhesion and motility (Fig. 8).

**Figure 8.**
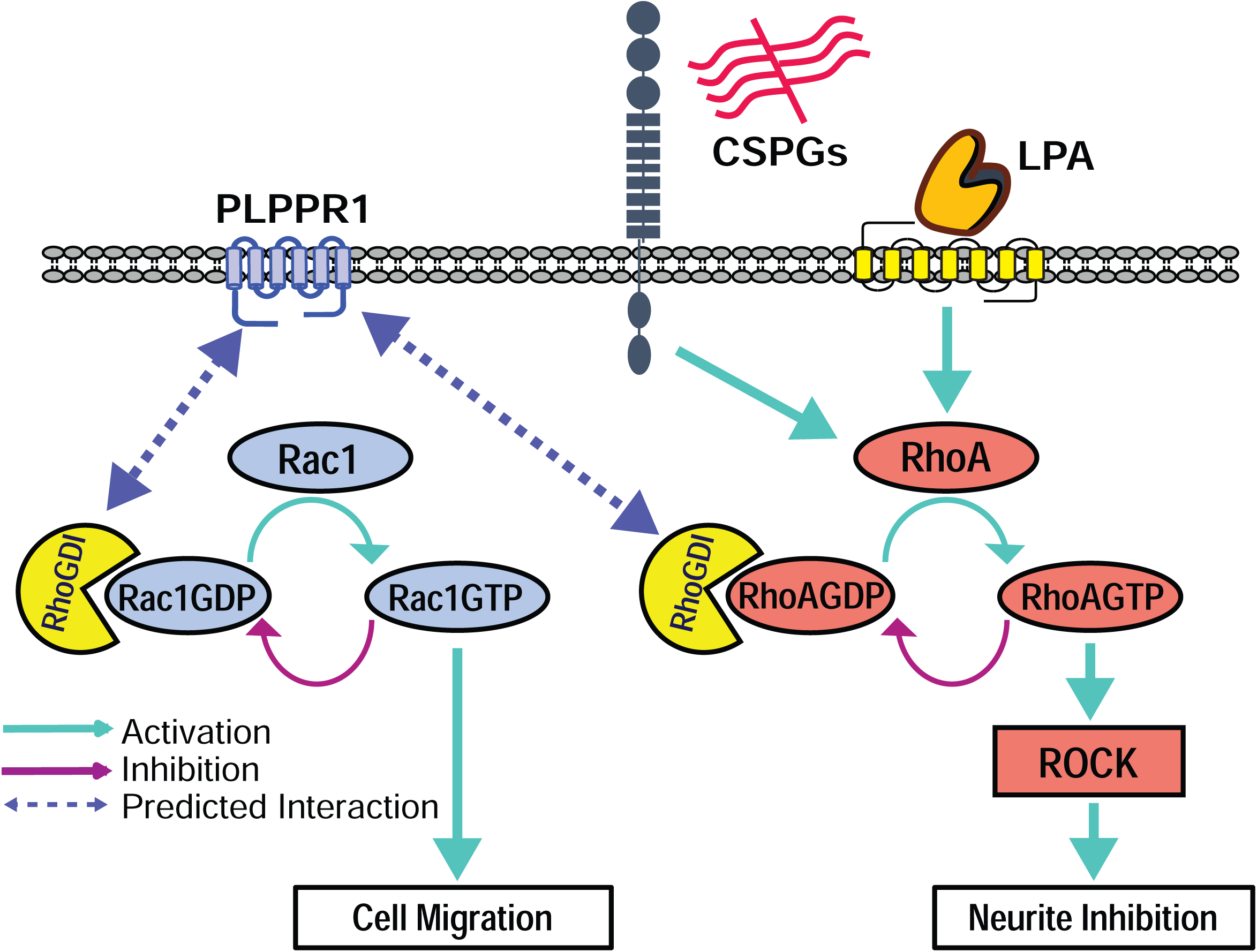
PLPPR1 modulates RhoA and Rac1 activity. Schematic showing the modulatory effect of PLPPR1 on RhoA and Rac1 activation through association with RhoGDI in response to CSPGs and LPA, to influence cell adhesion, cell migration and neurite outgrowth.

We first demonstrated that expression of PLPPR1 in Neuro2a cells results in decreased migration due to increased cell adhesion. In addition, migrating cells expressing PLPPR1 left long plasma membrane extensions that we have termed “trailing fibers”. Migrating cells often leave behind membrane fragments at the trailing edge of the cell (Regen and Horwitz, 1992), and their presence has been suggested to contribute to decreased cell advance (Chen, 1981). The persistence of these “trailing fibers” in cells expressing PLPPR1 suggest an inability of the cells to detach from the substrate during movement, likely due to the increase in adhesion to the fibronectin substrate.

Cell adhesion is mediated through the engagement and activation of integrins, which then promote the establishment of focal adhesions (Nobes and Hall, 1995; Rottner et al., 1999; Sinnett-Smith et al., 2001). Mechanical tension promotes the growth and maturation of focal adhesion complexes that is driven by RhoA-mediated myosin II activity (Pasapera et al., 2010). This is required for the coordinated propulsive force that enables cell migration (Gardel et al., 2010; Lauffenburger and Horwitz, 1996). PLPPR4 mediates cell adhesion through activation of ß1-integrins through the unique calmodulin-binding domain in its C-terminal tail (Liu et al., 2016). While PLPPR1 lacks the key motifs in its C-terminal tail, there is evidence that it may associate with β1-integrins (Yu et al., 2013). The nature of this interaction and its role in the observed increase in cell adhesion has yet to be explored.

Several studies have demonstrated that inhibition of Rho-ROCK activity prevents the maturation of focal adhesion complexes (Burridge and Guilluy, 2016; Sawada et al., 2002; Sinnett-Smith et al., 2001). Therefore, it is conceivable that the reduced RhoA activity and the nascent focal adhesion complexes that persist in cells expressing PLPPR1 may account for the increase in cell adhesion and decrease in cell migration. This is consistent with the increase in phosphorylation of paxillin and FAK observed in these cells, given that FAK mediated phosphorylation of paxillin plays a role in the assembly, turnover and stability of focal adhesions (Lawson et al., 2012; Pasapera et al., 2010; Turner, 2000; Webb et al., 2004; Zaidel-Bar et al., 2007). While increased phosphorylation of paxillin and FAK is observed in Neuro2a cells expressing PLPPR1, it remains unclear if this effect is due to direct interactions between PLPPR1 and focal adhesion complexes.

The PLPPR family of proteins share sequence and structural homology with the lipid phosphate phosphatase family (LPP), which regulate LPA signaling through phosphatase activity (Bräuer and Nitsch, 2008; Strauss and Bräuer, 2013). However, the PLPPR proteins lack key amino acid residues essential to the phosphatase activity of the LPP protein family and are not known to exhibit any phosphatase activity (Strauss and Bräuer, 2013). We show here that overexpression of PLPPR1 in hippocampal neurons overcomes neurite outgrowth inhibition induced by CSPGs. Both CSPGs and LPA are known to signal through activation of the RhoA GTPase (Alabed et al., 2006; Ohtake et al., 2016). Expression of PLPPR1 in Neuro2a cells reduced RhoA activation in response to LPA treatment. Furthermore, we confirmed the modulatory effect of PLPPR1 on RhoA activation of ROCK targets, MLC, MYPT1 and the ERM proteins. RhoA activation is controlled by regulatory proteins such as the GEFs and GDIs. GEFs catalyze the exchange of GDP for GTP, and GDIs sequester RhoA in the cytosol, regulating its translocation to the plasma membrane (Bokoch et al., 1994; Dovas and Couchman, 2005; Hodge and Ridley, 2016). RhoGDI was identified as a possible binding target for PLPPR1 in a proteomic screen for interacting proteins (Yu et al., 2013). Our results confirmed this interaction by co-immunoprecipitating PLPPR1 with RhoGDI. This association with RhoGDI was maintained in cells expressing the PLPPR1 C-terminal deletion mutant, PLPPR1ΔC43, suggesting an alternative site of association.

Without LPA stimulation, RhoA-GDP is tightly bound to RhoGDI (Boulter et al., 2010; Hodge and Ridley, 2016; Tkachenko et al., 2011). Upon LPA treatment, this interaction is disrupted, enabling the translocation of RhoA to the plasma membrane and its subsequent activation (Li et al., 2012). We have demonstrated here that expression of PLPPR1 maintains the interaction of RhoA with RhoGDI after LPA treatment suggesting that PLPPR1 may play a role in stabilizing the RhoA-RhoGDI complex, thereby, modulating the RhoA GDP exchange for GTP and its subsequent activation.

We show that expression of DN Rac1 exhibits a similar morphological change as expression of PLPPR1 in Neuro2a cells. Expression of DN Rac1 inhibits GTP loading by sequestering endogenous GEFs and forming non-functional RhoGTPase – RhoGEF complexes. This leads to decreased membrane ruffling and a failure to form proper lamellipodia (Debreceni et al., 2004; Ehrlich et al., 2002; McCarty et al., 2005). The co-expression of PLPPR1 with CA Rac1 rescues the trailing fiber morphology. CA Rac1 blocks the GAP binding domain and inhibits GTPase activity (Zhang et al., 2006). With an excess of mutated Rac1 fused to GTP, there may be an increased pool of GTP available to other RhoGTPases leading to increased RhoA activity in cells expressing CA Rac1 rescuing the morphology. Nucleation of actin filaments by Arp2/3 complex is driven by WAVE (WASP-family verprolin-homologous protein), downstream of Rac1 activation (Chen et al., 2017; Lai et al., 2008). Therefore, while Arp2/3 may not be required to induce membrane protrusions, localization of Arp2/3 is disrupted in cells overexpressing PLPPR1 and this may be a direct consequence of altered Rac1 activity.

Taken together, our results provide evidence for a novel signaling mechanism of PLPPR1– the modulation of RhoA and Rac1 activity by interaction with RhoGDI. This mechanism serves as a potential explanation for the ability of PLPPR1 to overcome CSPG, lipid and myelin-mediated inhibition of neurite outgrowth to promote axon regeneration after injury.

## MATERIALS AND METHODS

### Plasmid construction

pEGFP-PLPPR1 was prepared as described previously (Yu et al., 2015). Constitutively active Rac1 G12V (CA Rac1) and dominant negative Rac1 T17N (DN Rac1) were subcloned into XhoI/BamHI sites of pEGFP-C1 (Clontech, Mountain view, CA). mApple-FTR-940 (F-tractin) was originally from Michael Davidson’s laboratory (plasmid # 54902, Addgene, Cambridge, MA). mApple-paxillin was kindly provided by Dr. Clare Waterman (National Institutes of Health, Bethesda, MD). A construct expressing C-terminal HA-tagged PLPPR1 (PLPPR1–HA) was a generous gift from Dr. Andrew Morris (University of Kentucky, Lexington, KY) (Sigal et al., 2007).

### Cell culture and transfection

All animal procedures were performed in accordance with protocols approved by the Institutional Animal Care and Use Committee (IACUC) at the National Institutes of Health. Timed pregnant C57BL/6J mice were housed in a pathogen free facility with free access to food and water under a standard 12 h light/dark cycle.

Primary hippocampal neurons were isolated from embryonic day 18 – 20 C57BL/6J pregnant mice and transfected with 1 µg of either pEGFP or pEGFP-PLPPR1 per well using the 96-well shuttle Amaxa Nucleofector™ kit, program CA-138 (Lonza, Walkersville, MD). Neurons were grown in Neurobasal media supplemented with antibiotics (penicillin/streptomycin), glutamax, B27 (ThermoFisher Scientific, Waltham, MA) and 2 mM potassium chloride (KCl) for 48 h.

Neuro2a cells were maintained in DMEM supplemented with penicillin/streptomycin antibiotics and 10% fetal bovine serum (FBS). Twenty-four hours after plating, cells were transfected with either pEGFP or pEGFP-PLPPR1, using Avalanche Omni^®^ transfection reagent (EZ Biosystems, College Park, MD).

For CSPG treatment and neurite outgrowth analysis, transfected primary hippocampal neurons were plated onto glass coverslips coated with either poly-l-lysine (PLL) or a soluble CSPG mix containing neurocan, phosphacan, versican, and aggrecan isolated from embryonic chicken brain (CC117 Millipore, Billerica, MA). Neurons were subsequently immunostained for anti-βIII tubulin (Sigma Aldrich, St. Louis, MO) and imaged using a Nikon Eclipse Microscope with a 60×/1.4 NA oil immersion objective lens. Neurite outgrowth analysis was performed using the NeuronJ plugin of ImageJ.

All cell culture plates (Corning, Corning NY), 6-well (3506), 12-well (3512), 24-well (3527), 48-well (3548), MatTek dishes (35 mm petri dish with 14mm microwells, No. 1.5, MatTek Corporation, Ashland, MA) and Lab-Tek^TM^ culture II chamber slides, (155382) (ThermoFisher Scientific, Waltam, MA) were coated with 10 µg/ml fibronectin (Millipore, Billerica, MA), unless otherwise indicated, and incubated overnight at 4°C. Excess fibronectin was rinsed off by washing 3 times with phosphate buffered saline (PBS).

### Immunocytochemistry

Transfected Neuro2a cells were fixed in 4% paraformaldehyde (PFA), 48 h after plating and 24 h after transfection. Cells were briefly rinsed in PBS and incubated in blocking buffer (0.3% Triton X-100 in PBS, supplemented with 10% normal goat serum, NGS) for 1 h and then with primary antibody diluted in 0.3% Triton X-100 in PBS, supplemented with 2.5% NGS. Cells were washed in PBS and then incubated with secondary antibody diluted in 0.3% Triton X-100 in PBS, supplemented with 2.5% NGS. See Table 1 for dilutions and source of antibodies. Cells were then washed in PBS and then mounted with DAPI diluted in Fluoromount (Sigma-Aldrich). All phalloidin (ThermoFisher Scientific, Waltam, MA) staining was performed as previously described (Yu et al., 2015).

**Table 1.**
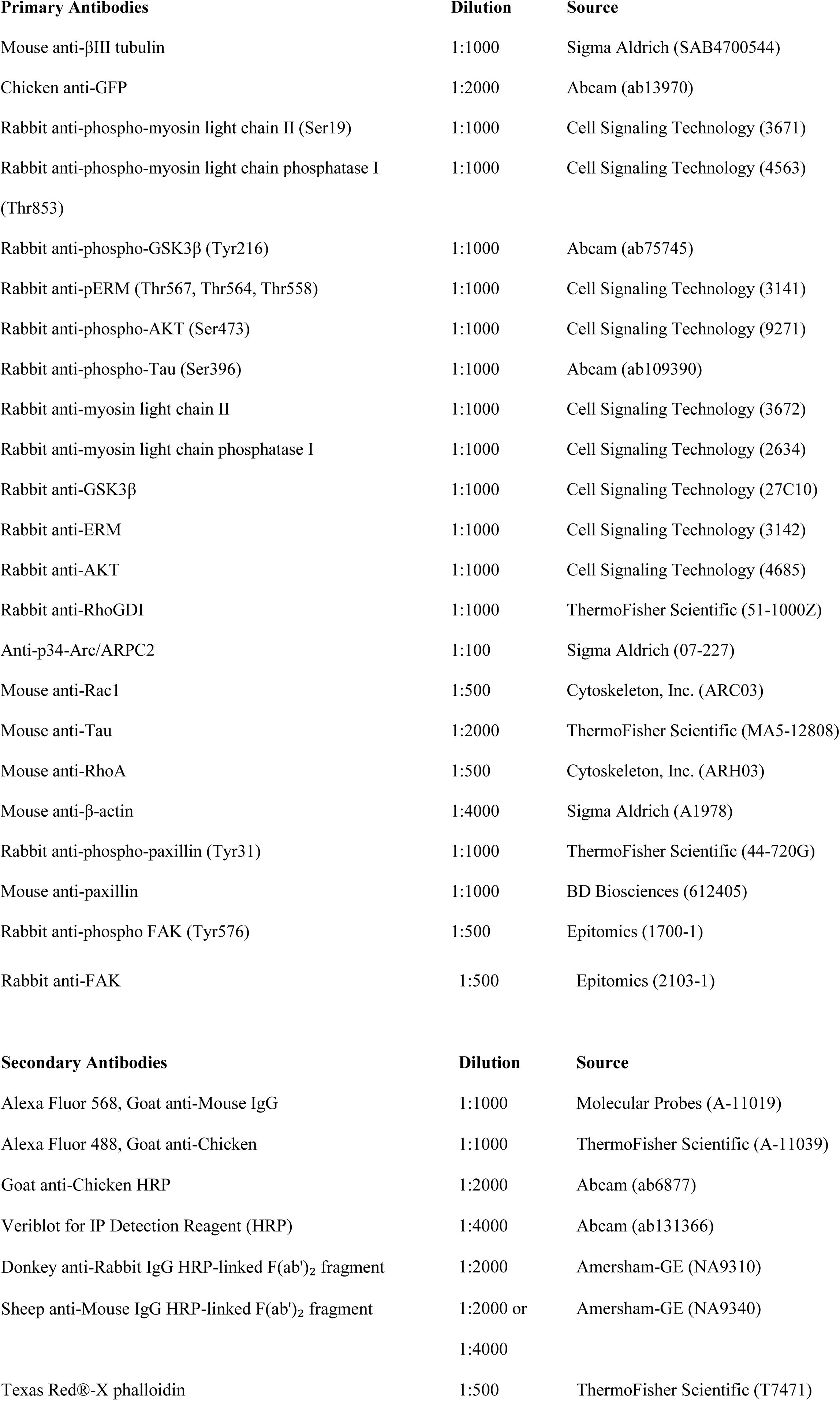
Antibodies Used.

| **Primary Antibodies** | **Dilution** | **Source** |
| --- | --- | --- |
| Mouse anti-βIII tubulin | 1:1000 | Sigma Aldrich (SAB4700544) |
| Chicken anti-GFP | 1:2000 | Abcam (ab13970) |
| Rabbit anti-phospho-myosin light chain II (Ser19) | 1:1000 | Cell Signaling Technology (3671) |
| Rabbit anti-phospho-myosin light chain phosphatase I (Thr853) | 1:1000 | Cell Signaling Technology (4563) |
| Rabbit anti-phospho-GSK3β (Tyr216) | 1:1000 | Abcam (ab75745) |
| Rabbit anti-pERM (Thr567, Thr564, Thr558) | 1:1000 | Cell Signaling Technology (3141) |
| Rabbit anti-phospho-AKT (Ser473) | 1:1000 | Cell Signaling Technology (9271) |
| Rabbit anti-phospho-Tau (Ser396) | 1:1000 | Abcam (ab109390) |
| Rabbit anti-myosin light chain II | 1:1000 | Cell Signaling Technology (3672) |
| Rabbit anti-myosin light chain phosphatase I | 1:1000 | Cell Signaling Technology (2634) |
| Rabbit anti-GSK3β | 1:1000 | Cell Signaling Technology (27C10) |
| Rabbit anti-ERM | 1:1000 | Cell Signaling Technology (3142) |
| Rabbit anti-AKT | 1:1000 | Cell Signaling Technology (4685) |
| Rabbit anti-RhoGDI | 1:1000 | ThermoFisher Scientific (51-1000Z) |
| Anti-p34-Arc/ARPC2 | 1:100 | Sigma Aldrich (07-227) |
| Mouse anti-Rac1 | 1:500 | Cytoskeleton, Inc. (ARC03) |
| Mouse anti-Tau | 1:2000 | ThermoFisher Scientific (MA5-12808) |
| Mouse anti-RhoA | 1:500 | Cytoskeleton, Inc. (ARH03) |
| Mouse anti-β-actin | 1:4000 | Sigma Aldrich (A1978) |
| Rabbit anti-phospho-paxillin (Tyr31) | 1:1000 | ThermoFisher Scientific (44-720G) |
| Mouse anti-paxillin | 1:1000 | BD Biosciences (612405) |
| Rabbit anti-phospho FAK (Tyr576) | 1:500 | Epitomics (1700-1) |
| Rabbit anti-FAK | 1:500 | Epitomics (2103-1) |
| **Secondary Antibodies** |  |  |
| Alexa Fluor 568, Goat anti-Mouse IgG | 1:1000 | Molecular Probes (A-11019) |
| Alexa Fluor 488, Goat anti-Chicken | 1:1000 | ThermoFisher Scientific (A-11039) |
| Goat anti-Chicken HRP | 1:2000 | Abcam (ab6877) |
| Veriblot for IP Detection Reagent (HRP) | 1:4000 | Abcam (ab131366) |
| Donkey anti-Rabbit IgG HRP-linked F(ab')<sub>2</sub> fragment | 1:2000 | Amersham-GE (NA9310) |
| Sheep anti-Mouse IgG HRP-linked F(ab')<sub>2</sub> fragment | 1:2000 or<br>1:4000 | Amersham-GE (NA9340) |
| Texas Red®-X phalloidin | 1:500 | ThermoFisher Scientific (T7471) |

For STED microscopy, cells were mounted in prolong diamond mounting media (ThermoFisher Scientific, Waltam, MA) and imaged on a Leica SP8 STED 3X/Confocal Microscope using 100×/1.4 oil immersion objective lens (Leica Microsystems, Buffalo Grove, IL). All other imaging was performed on a Zeiss 780 LSM confocal microscope using 63×/1.4 oil immersion objective lens.

### Live cell imaging

All live cell imaging was conducted on microscopes equipped with a heated stage at 37°C and 5% CO_2_ on either a Nikon A1R microscope (Nikon Instruments Inc., Melville, NY), a Zeiss 780 LSM confocal microscope (Carl Zeiss, Thornwood, NY) or an Delta Vision OMX microscope (GE, Marlborough, MA).

### Cell migration assay

Neuro2a cells transfected with either pEGFP or pEGFP-PLPPR1 were plated in Lab-Tek culture chamber slides coated with fibronectin. Time lapse images were acquired for 10 h at the rate of 1 frame/10 min on a Nikon A1R microscope using a 10×/0.45 NA objective lens. Cell migration was analyzed by tracking each transfected cell, using the middle of each cell body as the point of reference. The distance travelled was determined using NIS Elements software (Nikon Instruments Inc., Melville, NY).

### Cell adhesion assay

Neuro2a cells transfected with either pEGFP or pEGFP-PLPPR1 were plated in T75 culture flasks and detached with 5mM EDTA in PBS. Cells were re-plated in a 48-well plate coated with fibronectin at a density of 2.0 × 10^5^ cells per well. Cells were allowed to adhere for 1 h after plating and then washed briefly with DMEM culture media every 15 min thereafter. Fluorescence detection was performed on a SpectraMax M3 microplate reader (Molecular Devices, San Jose, CA). The cell density of adherent transfected cells was determined by fluorescence using the Softmax Pro7 software (Molecular Devices, San Jose, CA). Cell adhesion was calculated as the percentage of transfected cells that persists after each wash, from unwashed transfected cells. Cells were subsequently fixed in 4% PFA, rinsed twice in PBS and stained with DAPI. Images were acquired on a Nikon A1R microscope using a 10×/0.45 NA oil objective lens.

For the detachment assay, Neuro2a cells were plated on a 12-well plate coated with fibronectin and transfected 24 h after plating with either pEGFP or pEGFP-PLPPR1. After another 24 h, cells were briefly washed with DMEM culture media to remove serum and subsequently treated with Trypsin-EDTA diluted in DMEM (1:10) with increasing time intervals. Cells were then rinsed with PBS, fixed with 4% PFA and stained with DAPI. 10 × 10 tiled images were acquired on a Zeiss 780 LSM confocal microscope using a 10×/0.45 NA dry objective lens.

### Interference reflection microscopy

Neuro2a cells transfected with either pEGFP or pEGFP-PLPPR1 and 24 h post-transfection were fixed with 4% PFA for 15 min. Images were acquired with a Zeiss 780 LSM confocal microscope using a 63×/1.4 NA oil immersion objective lens. Single-frame TIFF images were extracted from TIFF stacks and processed using ImageJ. The area of each cell was traced from bright field images to obtain an ROI (region of interest). The image was then rescaled to an 8-bit scale of 0–255 based on pixel intensity values that fell within histogram bins exceeding 5% of the mean histogram bin height. A band pass filter was applied based on convolution with a 41 × 41 pixel kernel. The histogram was then stretched again based on a 5% cut-off of the mean values. A binary threshold was fixed by selecting pixel intensities above 90 on the 0 – 255 scale. The image was then inverted and the number of pixels above threshold was divided by the area of the ROI to determine intensity. A color map ‘Gem’ was used to display higher intensity pixels as blue and lower intensity as orange which represent stronger and weaker adhesions respectively. For quantification, the area of each cell was traced from bright field images to obtain the region of interest. A threshold was applied to the IRM image of the same cell and the number of pixels above threshold was divided by the ROI to calculate the area of cell attachment.

To measure the size of focal adhesion (FA) contacts, Neuro2a cells were plated on Fibronectin coated MatTek dishes at a density of 2 × 10^4^ cells and transfected with either pEGFP or pEGFP-PLPPR1. Cells were fixed and stained with anti-paxillin antibody. Images were acquired using TIRF microscopy on an inverted TIRF OMX Delta Vision microscope system using a 60×/1.49 NA TIRF objective lens. Counting of FAs in single cells (n = 10 cells) was done after noise removal by thresholding and applying a size constraint to FAs using ImageJ software.

### Actin turnover analyzed by FRAP

Neuro2a cells, plated on glass-bottom MatTek dishes coated with fibronectin, were co-transfected with F-tractin (mApple-FTR-940) and either pEGFP or pEGFP-PLPPR1. FRAP experiments were conducted on a Nikon A1R microscope using a 60×/1.4 NA oil objective lens. A ROI was selected from the leading edge of transfected cells and a 564 nm laser at 20% intensity was used as a pre-bleaching signal for F-tractin. The ROI was subsequently photobleached by three lasers at 458 nm, 564 nm and 633 nm, all set at 100% intensity. The post-bleaching acquisition was carried out at 60 frames at a rate of 1 frame/0.6 s. The fluorescence intensity was determined at every time point and automatically generated by NIS Elements software. Images were corrected for photobleaching and the fluorescence intensity of the ROI for each frame was normalized to the average initial intensity of pre-bleaching frames and to the area of the bleached cell. The fluorescence intensity for the pre-bleaching frames was calculated as the average fluorescence intensity in the first 5 frames, F_pre_. The value, F_0_, was designated as the average fluorescence intensity of the 8^th^ frame after the photobleaching phase that occurred in the 6^th^ and 7^th^ frames. Individual plots for each experiment was fitted to a model function. The fluorescence recovery of F-tractin was best fitted by a single-exponential function (double exponential function did not significantly improve the quality of the fit) using Matlab, revealing the presence of mobile actin fractions with different recovery kinetics. The following model was applied:

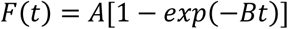

Where: A is the percentage of mobile fraction of actin and B is the time constant of fluorescence recovery. A and B were obtained from the fitted curve. The mean ± SEM was calculated for the percentage of fluorescence recovery and the time constant for each condition. The final graphs show the fluorescence recovery, mobile fraction of monomeric actin and t_1/2_ which indicates the time it took for F-actin polymerization.

### LPA treatment

Serum starved Neuro2a cells transfected with either pEGFP or pEGFP-PLPPR1 were treated with fatty-acid free bovine serum albumin, FAFBSA, (Sigma Aldrich, St. Louis, MO), or Lysophosphatidic acid, LPA (Cayman Chemical, Ann Arbor, MI) diluted in FAFBSA. Time lapse images were acquired on a Nikon A1R microscope with a 20×/0.75 dry objective lens with no time delay between frames for 20 min. Cell response was calculated as percentage of the total transfected cell count that retracted their neurites or underwent cell rounding.

### Immunoblot assay

Neuro2a cells seeded at a density of 4 × 10^5^ cells/well in a 6-well culture dish were transfected with either pEGFP or pEGFP-PLPPR1. Cells were either serum-starved for 16 – 18 h or maintained in DMEM supplemented with 10% FBS. Serum starved cells were then treated with either FAFBSA, 10% serum or increasing concentrations of LPA: 4, 8 and 16 μM for 2 min. Media was gently but completely removed from all cells and rinsed once with warmed PBS. Cell lysates were prepared in 2× SDS cell lysis buffer and clarified by centrifugation. Protein quantitation was performed using Ionic Detergent Compatibility Reagent for Pierce™ 660nm Protein Assay Reagent (ThermoFisher Scientific, Waltam, MA) or BCA method. Equal amount of proteins (20 µg/lane) was separated by SDS-PAGE and transferred onto a polyvinylidene difluoride (PVDF) membrane (Millipore, Billerica, MA). Membranes were blocked for 1 h in 5% nonfat dried milk in either 0.1% Tween20 in PBS (PBST) or 0.1% Tween20 in 10 mM Tris–HCl pH 7.5 (TBST) or blocked in 5% BSA in TBST. Membranes were incubated with primary antibodies overnight at 4°C, washed in either PBST or TBST and subsequently incubated with secondary antibody for 30 min. Antigen–antibody complexes were detected with KPL LumiGlo chemiluminescent substrate (Seracare, Milford, MA). See Table 1 for source and dilutions for all primary and secondary antibodies used.

### Co-immunoprecipitation assay

Neuro2a cells were seeded at a density of 4 × 10^5^ cells/well in a 6-well culture dish and transfected with either pEGFP or pEGFP-PLPPR1. Cells were harvested, washed twice in ice-cold PBS and then resuspended in ice-cold cell lysis buffer (10 mM Tris pH 7.5, 150 mM NaCl, 0.5 mM EDTA, 1% NP-40 supplemented with protease inhibitor cocktail set V EDTA free (Millipore, Billerica, MA) and phosphatase inhibitors (Thermo Fisher Scientific, Waltham, MA). Cell lysates were cleared and then incubated with GFP-Trap MA beads as instructed in ChromoTek GFP-Trap^®^ MA immunoprecipitation kit (ChromoTek Inc., Hauppauge, NY). Cell lysates were subsequently washed twice in dilution buffer (10 mM Tris-HCl pH 7.5, 150 mM NaCl, 0.5 mM EDTA, 0.5% NP-40 supplemented with protease and phosphatase inhibitors) and then once in wash buffer (10 mM Tris-HCl pH 7.5, 500 mM NaCl, 0.5 mM EDTA, 0.5% NP-40 supplemented with protease and phosphatase inhibitors). Proteins were eluted and subjected to SDS-PAGE as described above.

To immunoprecipitate RhoGDI, transfected cells were rinsed in warmed PBS, harvested in cell lysis buffer (20 mM Tris pH 7.5, 150 mM NaCl, 0.5 mM EDTA, 1% NP-40 supplemented with protease and phosphatase inhibitors) and precleared with rabbit IgG serum and Protein G agarose beads (Seracare, Milford, MA). Samples were subsequently incubated with anti-RhoGDI overnight at 4°C and then incubated with Protein G agarose beads for 1 h at 4°C. Samples were washed twice in dilution buffer (20 mM Tris pH 7.5, 150 mM NaCl, 0.5 mM EDTA, 0.5% NP-40 supplemented with protease and phosphatase inhibitors) and once in wash buffer (20 mM Tris pH 7.5, 500 mM NaCl, 0.5 mM EDTA, 0.5% NP-40 supplemented with protease and phosphatase inhibitors). Proteins were eluted and subjected to SDS-PAGE as described above.

### RhoA and Rac1 pull-down assay

Serum starved Neuro2a cells transfected with either pEGFP or pEGFP-PLPPR1 were treated with either FAFBSA, 10% serum or 16 µM LPA. Briefly, cells were rinsed with warmed PBS and cell lysates prepared in cell lysis buffer as instructed and provided by the RhoA or Rac1 pull-down instruction manual and kit (Cytoskeleton, Inc. Denver, CO). Protein quantitation was performed by BCA method and 600 μg of protein was incubated with 50 μg Rhotekin-RBD beads for RhoA pull-down or with 10 µg of PAK-PBD beads for Rac1 pull-down at 4°C for 1 h. Complexes were washed in wash buffer provided in the pull-down kit and protein was eluted in 2× SDS sample buffer and SDS-PAGE performed as described above.

### Statistical analysis

All analyses were performed using GraphPad Prism software version 7 (GraphPad Software, La Jolla, CA). Data was analyzed by unpaired Student’s *t* tests, one-way ANOVA or two-way ANOVA with Welch’s correction or Tukey’s post-hoc multiple comparison test to determine significance. Normality of the distribution of the data was tested with Kolmogorov–Smirnov normality tests using the column statistics function of GraphPad Software. All tests were two-tailed with significance indicated as follows: *p<0.05; **p<0.01; ***p<0.001; ****p<0.0001. Unless otherwise specified, values represent the means ± SEM. All experiments were repeated independently at least three times.

## Online supplementary material

Fig. S1 shows cells overexpressing PLPPR1 resist trypsin detachment from the fibronectin substrate. Fig. S2 shows that expression of PLPPR1 has no effect on phosphorylated levels of GSK3ß, Tau or AKT after LPA treatment. Fig. S3 shows that without LPA stimulation, PLPPR1 has no effect on ROCK targets, MLC, MYPT1, pERM, including GSK3ß, Tau and AKT.

## Acknowledgements

We wish to thank Daniela Malide and Xufeng Wu of the NHLBI Light Microscopy Core facility for their help and guidance and Clare Waterman for materials provided.

This work was funded by the NIH Intramural Research Program. The authors declare no competing financial interests.

Author contributions: C.A. Iweka, S. Tilve and H.M. Geller conceived the project. C.A. Iweka and S. Tilve performed the experiments. C.A. Iweka, S. Tilve, C. Mencio and H.M. Geller wrote the manuscript. All authors planned the experiments, analyzed and discussed the results and commented on the manuscript.

## Abbreviations

PLPPR: Phospholipid Phosphatase-Related Protein
FA: Focal Adhesions
CSPG: Chondroitin Sulfate Proteoglycan
PLL: Poly-L-Lysine
LPA: Lysophosphatidic acid
LPP: Lipid Phosphate Phosphatase
FAFBSA: Fatty Acid Free Bovine Serum Albumin
MLC: myosin light chain II
MYPT1: myosin light chain phosphatase 1
ERM: Ezrin, Radixin and Moesin
GSK: Glycogen Synthase Kinase
GEF: Guanine Exchange Factor
GAP: GTPase Activating Protein
RhoGDI: Rho Guanine Nucleotide Dissociation Inhibitor
IRM: Interference Reflection Microscopy
TIRF: Total Internal Reflection Fluorescence
STED: STimulated Emission Depletion
FRAP: Fluorescence Recovery After Photobleaching
ROCK: Rho-associated protein kinase
CA: Constitutively Active
DN: Dominant Negative
FAK: Focal adhesion kinase
GTP: Guanosine triphosphate
GDP: Guanosine diphosphate
WAVE: WASP-family verprolin-homologous protein).

---

**Movie S1. “Trailing fibers” observed in migrating PLPPR1 cells with Interference Reflection Microscopy.** Time lapse images of Neuro2a cells transfected with either pEGFP or pEGFP-PLPPR1 were acquired in bright field using Interference Reflection Microscopy (IRM) on Zeiss 780 LSM confocal microscope system using a 10×/0.45 NA objective lens. Movie shows 3 min/frame.

**Movie S2. Focal adhesions remain nascent in cells overexpressing PLPPR1 compared to cells expressing EGFP.** Time lapse images of Neuro2a cells co-expressing mCherry-paxillin with either EGFP (left) or EGFP-PLPPR1 (right) were acquired on Delta Vision OMX microscope system using a 60×/1.49 NA TIRF objective lens. Movie highlight the size and nature of focal adhesions, including formation of trailing fibers in cells co-expressing mCherry-paxillin with EGFP-PLPPR1. Movie shows 3 min/frame.

**Movie S3. LPA induces neurite retraction and cell rounding in Neuro2a cells.** Serum starved Neuro2a cells were exposed to either FAFBSA (left) or 16 µM LPA (right). Time lapse images were acquired on a Nikon A1R microscope with a 20×/0.75 dry objective lens with no time delay between frames for 20 min. Movie shows 1 min/frame.

**Movie S4. PLPPR1 impedes LPA induced neurite retraction in Neuro2a cells.** Serum starved Neuro2a cells transfected with either pEGFP (left) or pEGFP-PLPPR1 (right) were exposed to 16 µM LPA. Time lapse images were acquired on a Nikon A1R microscope with a 20×/0.75 dry objective lens with no time delay between frames for 20 min. Movie shows 1 min/frame.

**Figure S1.**
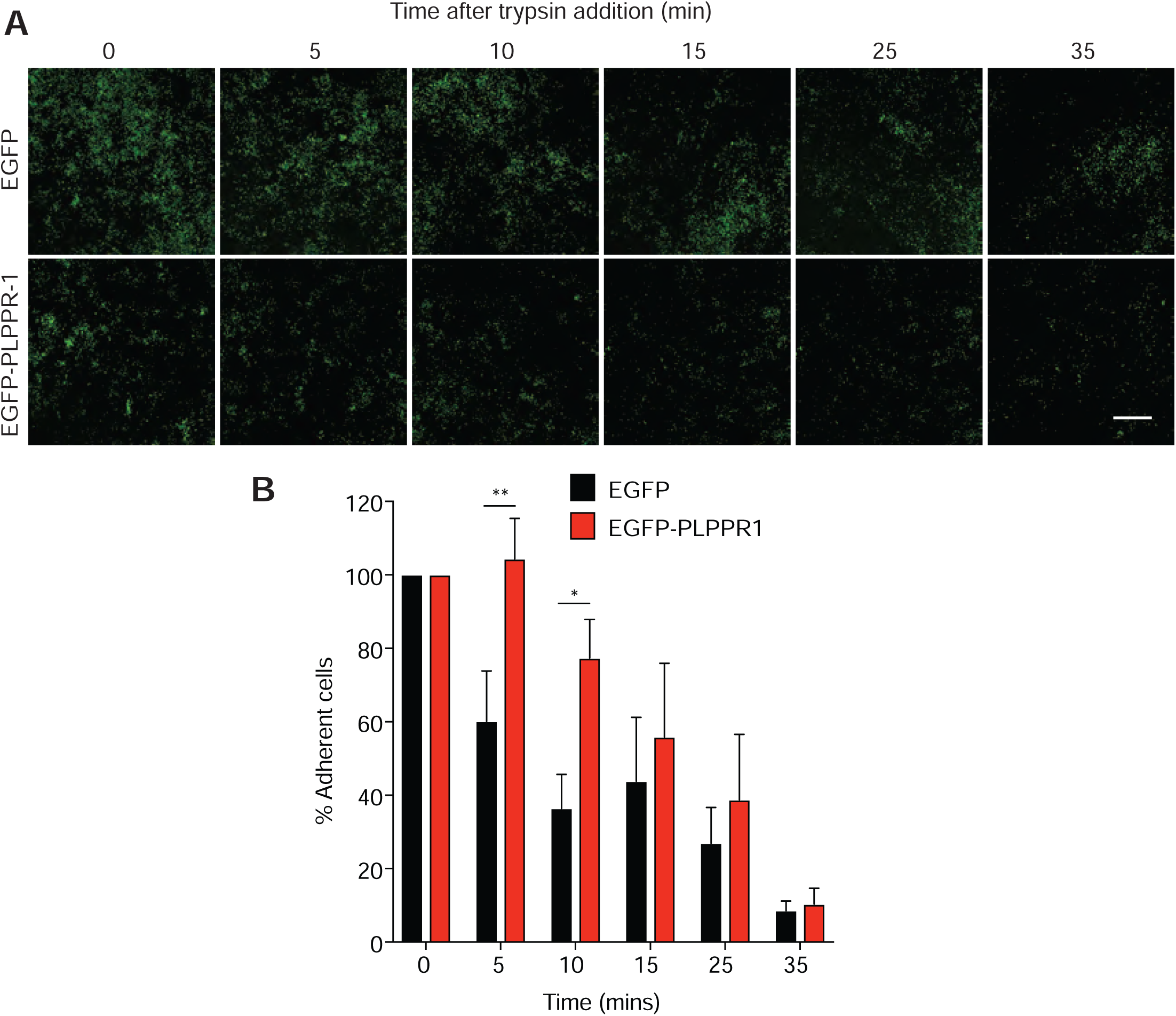
Cells expressing PLPPR1 are resistant to detachment. **(A)** Representative images of Neuro2a cells transfected either pEGFP (top) or pEGFP-PLPPR1 (bottom) and subjected to trypsin/EDTA treatment over time. Scale bar, 100µm. **(B)** Quantification of the number of adherent cells resistant to detachment was determined as a percentage of total unwashed transfected cells. Data represent mean ± SEM. p-values were calculated using Two-way ANOVA repeated measures with Tukey posthoc analysis. *p<0.05, **p<0.01. Experiment was conducted in triplicate.

**Figure S2.**
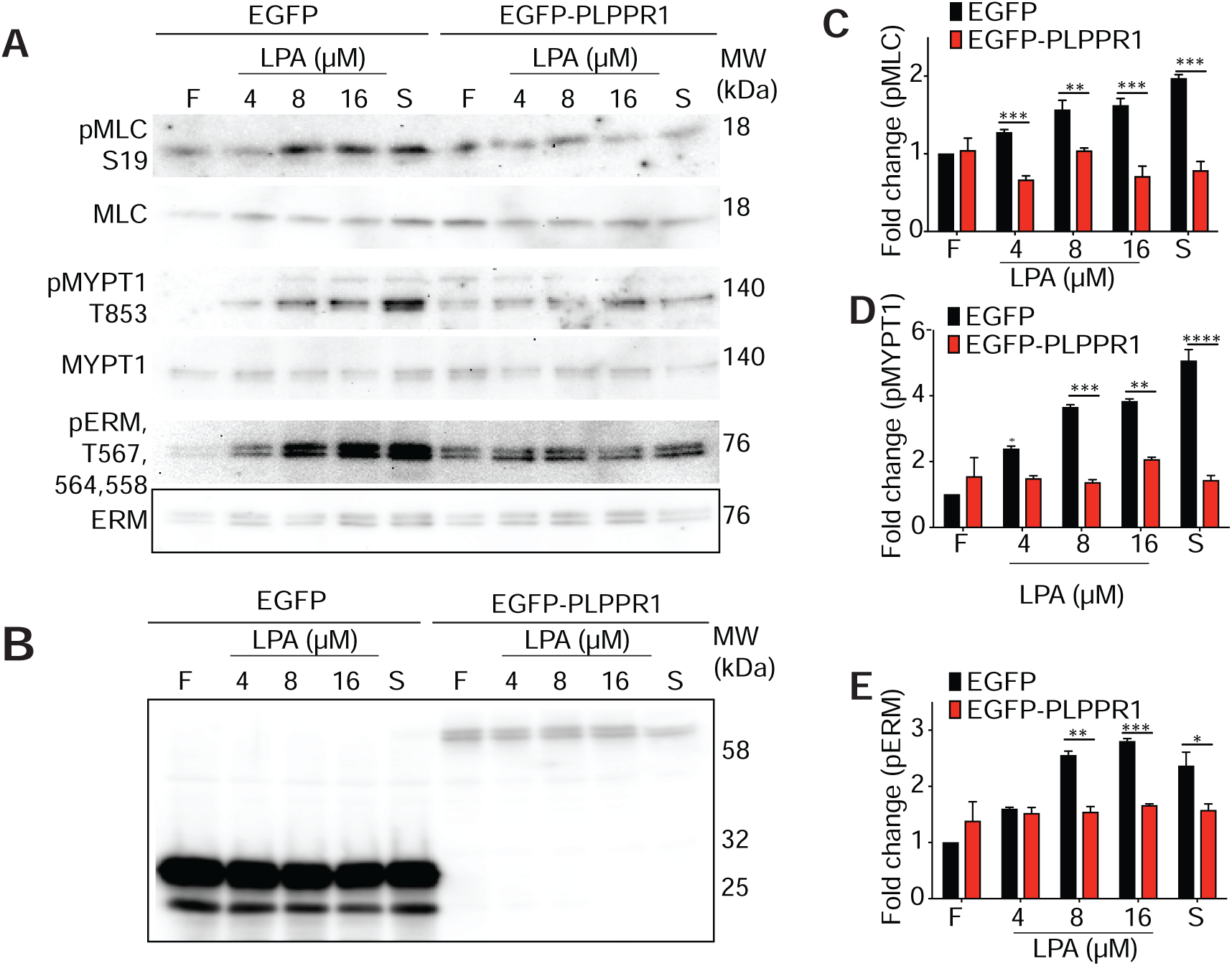
Expression of PLPPR1 reduces phosphorylation levels of RhoA downstream effectors after LPA treatment. Neuro2a cells transfected with either pEGFP or pEGFP-PLPPR1 were serum starved and treated with either FAFBSA (F), serum (S), or LPA for 5min. **(A)** Immunoblots of cells transfected with either pEGFP or pEGFP-PLPPR1 and treated with FAFBSA, serum, or LPA were probed for pMLC and MLC, pMYPT1 and MYPT1, pERM and ERM. **(B)** Immunoblot for EGFP to show equal expression. Densitometry analysis of phosphorylated **(C)** MLC, **(D)** MYPT1, and **(E)** ERM versus total protein was performed using Image Studio Lite. Data represents mean ± SEM. p-values were calculated using Two-way ANOVA with Tukey’s posthoc analysis, *p<0.05, **p<0.01, ***p<0.001, ****p<0.0001 respectively. All experiments were conducted in triplicate.

**Figure S3.**
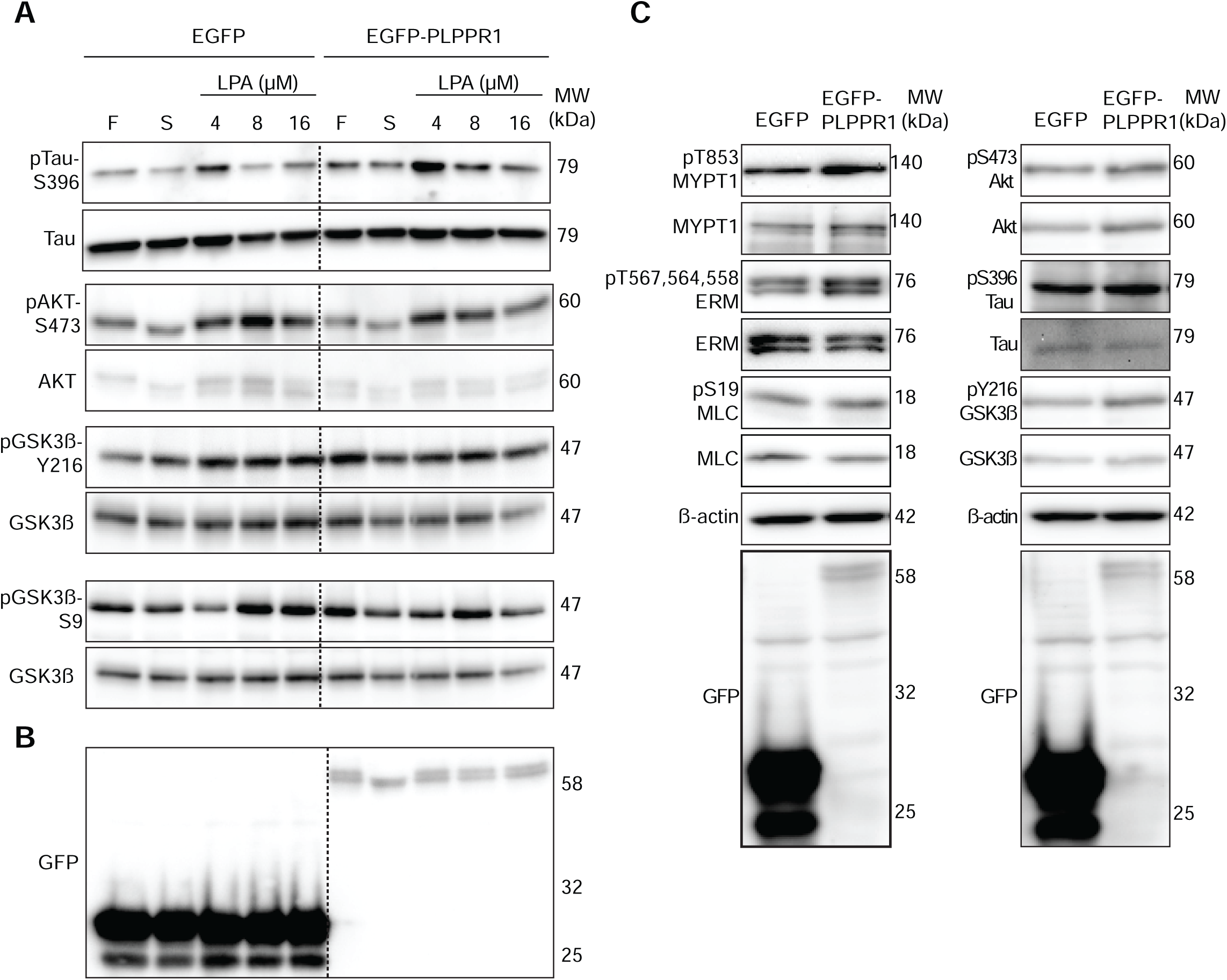
Effect of PLPPR1 expression on non-ROCK target phosphorylation after LPA treatment and all targets without LPA. **(A)** Immunoblots showing phosphorylated levels of several ROCK targets Tau, AKT, and GSK3ß in serum starved Neuro2a cells expressing EGFP or EGFP-PLPPR1 and exposed to FAFBSA (F), LPA or serum (S) for 5 min. **(B)** All membranes were stripped and reprobed for each total protein and GFP. **C)** Immunoblots showing phosphorylated levels of several ROCK targets, MYPT1, ERM, MLC, AKT, Tau, GSK3ß in Neuro2a cells expressing EGFP or EGFP-PLPPR1. All membranes were stripped and reprobed for each total protein and GFP. ß-actin was used as loading control.

